# *Saccharomyces cerevisiae* gene expression during fermentation of Pinot noir wines at industrially relevant scale

**DOI:** 10.1101/2021.01.11.426308

**Authors:** Taylor Reiter, Rachel Montpetit, Shelby Byer, Isadora Frias, Esmeralda Leon, Robert Viano, Michael Mcloughlin, Thomas Halligan, Desmon Hernandez, Ron Runnebaum, Ben Montpetit

## Abstract

During a wine fermentation, *Saccharomyces cerevisiae* transforms grape must through metabolic activities that generate ethanol and other compounds. Thousands of genes change expression over the course of a wine fermentation to allow *S. cerevisiae* to adapt to and dominate the fermentation environment. Investigations into these gene expression patterns have previously revealed genes that underlie cellular adaptation to the grape must and wine environment involving metabolic specialization and ethanol tolerance. However, the vast majority of studies detailing gene expression patterns have occurred in controlled environments that do not recapitulate the biological and chemical complexity of fermentations performed at production scale. Here, we present an analysis of the *S. cerevisiae* RC212 gene expression program across 40 pilot-scale fermentations (150 liters) using Pinot noir grapes from 10 California vineyards across two vintages. We observe a core gene expression program across all fermentations irrespective of vintage similar to that of laboratory fermentations, in addition to novel gene expression patterns likely related to the presence of non-*Saccharomyces* microorganisms and oxygen availability during fermentation. These gene expression patterns, both common and diverse, provide insight into *Saccharomyces cerevisiae* biology critical to fermentation outcomes at industry-relevant scales.

**Importance:** This study characterized *Saccharomyces cerevisiae* RC212 gene expression during Pinot noir fermentation at pilot scale (150 liters) using production-relevant conditions. The reported gene expression patterns of RC212 is generally similar to that observed in laboratory fermentation conditions, but also contains gene expression signatures related to yeast-environment interactions found in a production setting (e.g., presence of non-*Saccharomyces* microorganisms). Key genes and pathways highlighted by this work remain under-characterized, raising the need for further research to understand the roles of these genes and their impact on industrial wine fermentation outcomes.

## Introduction

*Saccharomyces cerevisiae* is most often the dominant fermentative organism during vinification. As a domesticated species, it has evolved specialized metabolic strategies to assimilate sugars in grape must and transform them into ethanol, thereby outcompeting other microorganisms during fermentation (1). During this process, *S. cerevisiae* encounters a dynamic stress landscape. In early fermentation, sources of stress include high sugar concentration (osmotic stress), low pH (acid stress), decreasing oxygen (hypoxia), the presence of other organisms that compete for nutrients or produce inhibitory compounds, and sulfur dioxide additions that are used to inhibit spoilage organisms. As fermentation progresses, nutrients become limiting (starvation), temperature may rise or be kept low (heat/cold stress), and ethanol concentrations rise (ethanol stress). Yet, through a coordinated gene expression response, *S. cerevisiae* adapts to these stresses and most often continues fermentation until the must is dry.

High throughput gene expression profiling (e.g., microarray and RNA sequencing) has offered a window into the metabolic strategies used by *S. cerevisiae* during fermentation to adapt and dominate fermentation environments. Previous research has reported expression changes in >2000 genes during fermentation (2–4). In early fermentation, this is marked by expression of gene products that support biosynthetic processes and acquisition of abundant nutrient resources (2, 3). As fermentation progresses, nitrogen limitation, phosphate limitation, and/or ethanol accumulation can trigger a transition to a non-proliferative state (i.e., stationary phase), which involves remodeling the gene expression program to support cellular adaptation to the changing environmental with continued metabolism (2, 3). Towards the end of fermentation, relief of nitrogen catabolite repression (2) and increased expression of nitrogen recycling genes (2, 5) is observed, which can be accompanied by further remodeling of the translational machinery and increased oxidative metabolism (5, 6). As ethanol concentrations rise through the end of fermentation, a gradual transcriptome response to ethanol stress is also observed (3). This response overlaps with, but appears distinct from, the environmental stress response seen in laboratory yeast (2, 3, 7), which may be related to the multitude of simultaneous stresses experienced by the yeast at the end of a wine fermentation (Bisson 2019). Indeed, the majority of genes with stress response elements in their promoter are expressed at the end of fermentation (8).

Through the associated metabolic processes that consume and produce a multitude of compounds, *S. cerevisiae* gene expression in response to environmental factors is related to overall fermentation kinetics and wine sensory outcomes. For example, fermentations can become sluggish or stuck when *S. cerevisiae* inadequately adapts to stresses encountered in the wine fermentation environment (9). In addition, altered gene expression likely underlies differential wine sensory characteristics in fermentations conducted with different industrial yeast strains (10, 11). To impact wine quality, genetic strategies have been applied in an attempt to alter the expression of flavor-associated genes (12), which have achieved variable levels of success. Consequently, further study of the *S. cerevisiae* gene expression program across fermentation is required to understand the yeast-environment relationship and how these interactions may be controlled to alter fermentation outcomes.

Given the importance of the yeast-environment interaction in determining gene expression, a major consideration with respect to collecting such data is the fermentation conditions used. To date, the majority of gene expression surveys have profiled fermentations that deviate in one or more ways from the industrial conditions in which most fermentations take place. For example, hundreds to thousands of liters of grape must are fermented to wine at industrial scales, while milliliter to liter volumes are commonly used in laboratory studies of gene expression (2–5, 13–16). Industry scale fermentations also have different kinetics than lab scale fermentations (4, 15, 17), and are less aromatic due to differences in hydrodynamics (15, 18). Similarly, dissolved oxygen differs at lab scale from industrial scale (4), which can impact fermentation outcomes (19). Possibly reflecting these different environments, at the end of fermentation the expression of key genes involved in amino acid transport and other core metabolic processes have been shown to differ between lab and industrial fermentations (4). Consequently, we propose that the physical and chemical differences in lab versus industrial scale wine fermentations are important factors to consider when analyzing gene expression patterns across fermentation.

Another major consideration when conducting gene expression studies, is that most studies investigate the fermentative capability of *S. cerevisiae* in monoculture using sterile synthetic media or filter sterilized grape must (2–5, 12). These controlled studies are important and allow connections between the media, gene expression, and wine outcomes to be made (12), but do not recapitulate the complexity of a natural grape must that varies in parameters like nitrogen composition, pH, and phenolic and elemental profiles (20–23). In addition, these experiments lack the diverse grape must microbiome that is a contributing component of wine fermentations (24–38). These are all parameters that will shape the fermentation environment and the metabolic response of *S. cerevisiae*.

Inter-species interactions are a critical component of the fermentation environment that informs the biology and behavior of *S. cerevisiae* during fermentation. It has been shown that non-*Saccharomyces* yeast impact the metabolism of *S. cerevisiae* through direct and indirect interactions (39–41), leading to faster resource acquisition by *S. cerevisiae* in early fermentation and altered metabolism of vitamins and minerals (40–43). While research is still needed to describe the impact of a diverse microbial consortia on *S. cerevisiae* during fermentation (44, 45), it remains that industrial fermentations are not sterile and involve diverse microorganisms (28, 34, 35, 37, 38). Even in fermentations treated with sulfur dioxide (SO*_2_*) to control microbial spoilage organisms, native fungi and bacteria are metabolically active during fermentation (38, 46, 47). This makes profiling *S. cerevisiae* gene expression amongst diverse microbial consortia important, as it will lead to a better understanding of the principles that govern *S. cerevisiae* gene expression and metabolism during fermentation.

Here, to begin to address the impact of an industrial wine fermentation environment on *S. cerevisiae* gene expression, we incorporate the inherent variability found in industrial fermentations and determine the *S. cerevisiae* RC212 gene expression program across chemically and biologically diverse Pinot noir grape musts. Specifically, time series RNA-sequencing was used to capture the gene expression profiles of RC212 during 40 inoculated primary fermentations at pilot scale (150 liters) using California Pinot noir grapes from 10 vineyards across two vintages. Using this data, a core metabolic program was defined during fermentation, which is well reflected by lab-scale fermentations, in addition to gene expression patterns that deviate from expectation. In particular, we observe altered gene expression that may be explained by the presence non-*Saccharomyces* organisms and regulation of metabolic processes related to stress, oxygen, and redox balance throughout fermentation. These observations suggest that the core genetic programs uncovered by lab-based studies are detected in industry-relevant fermentations, but production-based environmental factors induce other gene expression programs that are layered on top of the core gene expression program. We expect that understanding such variations in gene expression within a wine production-like environment will be key to defining approaches that can be used to manage commercial fermentation outcomes.

## Results & Discussion

### Conditions and rates of fermentation

Pinot noir grapes were harvested from the same 10 vineyards in California during the 2017 and 2019 vintages for wine production at the UC Davis Teaching & Research Winery (**Figure 1A**). To standardize fermentations, grapes from the same Pinot noir clone and rootstock were harvested at the same ripeness (∼24 Brix, total soluble solids as a proxy for sugar concentration). We sampled duplicate fermentations that used the grape material from each vineyard for a total of 40 fermentations (20 from each vintage) at industry-relevant scales using the same wine making protocol. Each fermentation was inoculated with the commercial wine strain *S. cerevisiae* RC212 and sampled to collect cells for gene expression analysis at 16 (exponential phase / early fermentation), 64 (stationary phase / mid fermentation), and 112 (decline phase / end of fermentation) hours post-inoculation (**Figure 1B**). While sampling times were standardized across fermentations, the rate of fermentation varied, resulting in samples being collected across a range of Brix values (**Figure 1C**). Differences in fermentation rates likely reflect diversity in the starting material and differential fermentation outcomes, which has also been demonstrated in sensory studies performed on wines produced from these vineyard sites in previous vintages (48).

**Figure 1:**
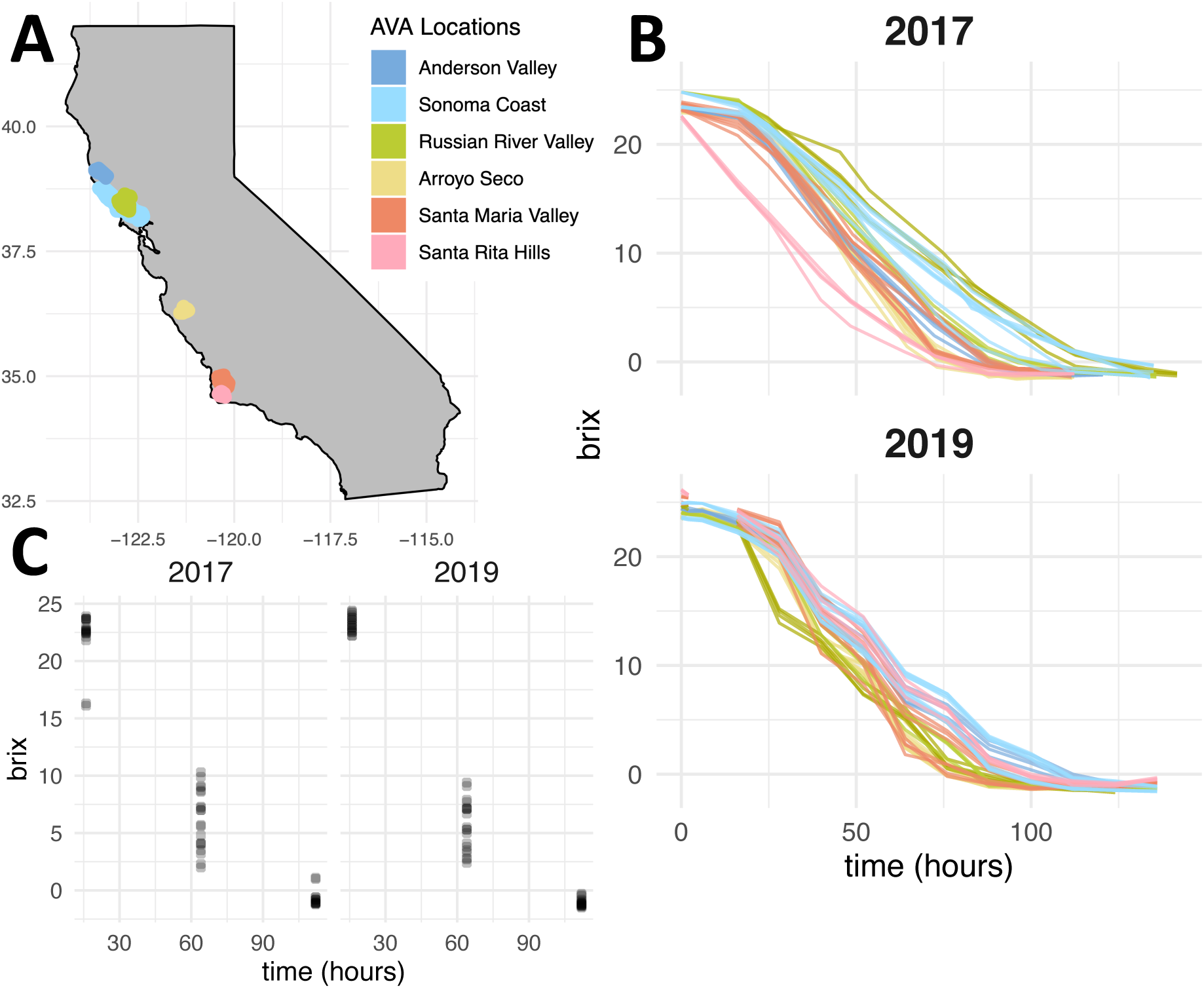
California vineyard locations and fermentation patterns. **A).** Map displaying the six American Viticulture Areas (AVAs) in which the 10 study vineyards are located. **B.)** Fermentation curves reflecting the change in Brix over fermentation. Brix is a measure of total soluble solids that is used as a proxy for sugar concentration in grapes, grape must, and wine. **C).** Brix at time of sampling for each RNA-seq sample relative to inoculation. While samples were taken at the same absolute time, fermentations proceeded at different rates leading to different Brix values in each fermentation.

### A consistent whole-transcriptome remodeling occurs during fermentation independent of vintage

Using 3′ Tag RNA sequencing (3′ Tag-seq), we profiled *S. cerevisiae* RC212 gene expression at 16, 64, and 112 hours after inoculation from both the 2017 and 2019 vintages. Since these samples provided gene expression measurements across a multitude of Brix values (**Figure 1C**), data from all 10 sites was combined and used to perform differential expression along the continuous variable Brix. The resulting data defines a core vineyard-independent gene expression program of RC212 during California Pinot noir fermentations. Under this model, log*_2_* fold change values represent the change in gene expression for each unit increase of Brix. Given that Brix decreases during fermentation, a positive log*_2_* fold change corresponds to a gene that decreased in expression as fermentation progressed, while a negative log*_2_* fold value corresponds to a gene that increased in expression as fermentation progressed (see examples in **Figure 2A**). After performing differential expression, we further intersected the differentially expressed genes across vintages to determine consistent changes that were vintage-independent. From this analysis, 991 genes decreased expression as Brix decreased, while 951 genes increased expression as fermentation progressed (**Figure 2AB, Table S1**). Each vintage also showed unique differential gene expression patterns, which may occur due to vintage-specific differences in fermentation. However, we generated these data at different times and applied newly developed methods (UMI barcoding, see methods) for sequencing the 2019 samples, and as such we suspect that the higher number of differentially expressed genes in the 2017 vintage may reflect differences in sequencing data quality. Nonetheless, the large fraction of shared differentially expressed genes suggests that a core gene expression program is followed independent of vintage.

**Figure 2:**
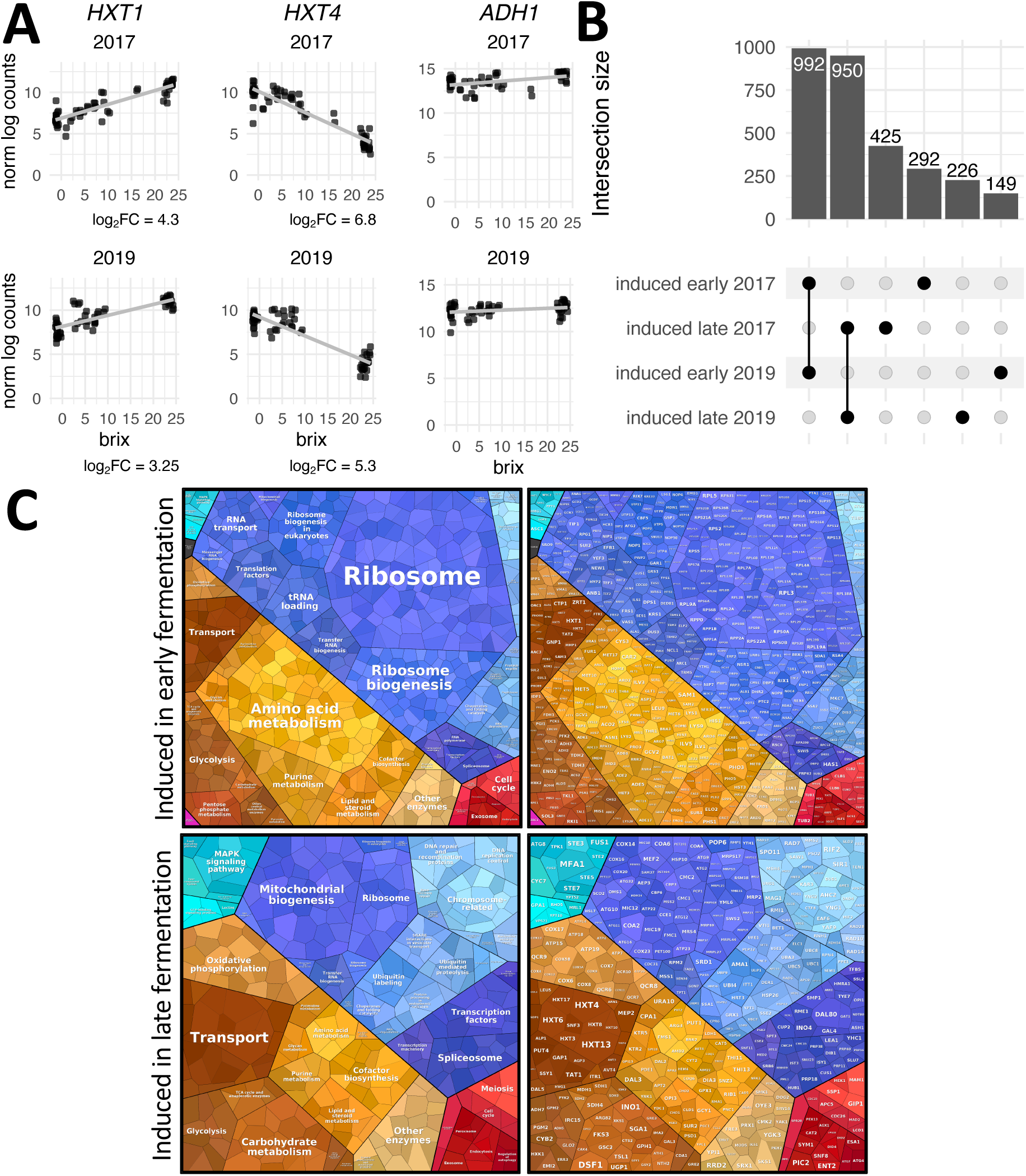
Transcriptome remodeling in fermentation is consistent across fermentations and vintages. **A.)** Genes expressed in early (*HXT1*), late (*HXT4*), and constitutively (*ADH1*) in fermentation. **B.)** Upset plot showing the intersection of genes that are significantly expressed at the beginning and end of fermentation in each vintage using a log_2_ fold change cut off of 1. The majority of genes are consistently expressed across fermentations and vintages. **C.)** Proteomaps depicting Gene Ontologies (left) and genes (right) that are expressed in early (top) and late (bottom) fermentation. The size of individual genes reflects the associated log_2_ fold change value. Note that for presentation purposes not all genes that are significantly expressed are depicted. See Table S1 for a complete list of genes.

Of the genes differentially expressed in fermentation and shared across vintage, many are known to function in wine fermentation and are central to yeast growth, metabolism, and cell survival (**Figure 2B and C**, **Table 1**). A strong signature of growth early in fermentation is observed that included cellular investment in ribosome biogenesis, metabolism of lipids, purines, and amino acids, as well as cell division machinery (**Figures 2C and S1-S2**). These processes, coupled with enrichment of associated pathways involved in RNA transcription and transport, reflect energy use for cell growth and proliferation associated with log phase growth occurring in early fermentation. Further in fermentation, changes in ribosomal machinery gene expression occurred, as reported in previous studies (49) (**Figure 2C**), reflecting a transition to a non-proliferative metabolic state. Late in fermentation this was accompanied by changes in gene expression linked to nutrient limitation, altered metabolism, and entry into meiosis (**Figures 2C and S3-S4**), which included gene expression patterns consistent with hallmark isoform switches in hexose transporters and glycolytic enzymes that occur as concentrations of glucose or fructose change (50) (**Figures 2B and 2C**). For example, *HXT1* encodes a low affinity glucose transporter that was more strongly expressed at the beginning of fermentation when glucose is abundant. *HXT4* has a high affinity for glucose and is expressed when glucose concentrations are low (51), which is also observed in our data as *HXT4* expression increased in late fermentation. Importantly, the pathways we have identified as enriched in early and late fermentation align with expectations based on previous research and the known biology of *S. cerevisiae* during fermentation (2–5, 13–16). This highlights the core processes that previous research efforts have defined and provides confidence that the analysis methods employed in these pilot-scale fermentations capture these biologically important transitions.

**Table 1:** Genes differentially expressed throughout fermentation shared across vintages

| Wine process | Gene | Expressed most highly | Gene product function |
| --- | --- | --- | --- |
| Glycolysis and fermentation | *HXT3* | late | Hexose transporter induced by both high and low glucose concentrations. |
|  | *PFK1, PFK2* | *PFK2*: early | Phosphofructokinases that catalyze the first irreversible reaction specific to glycolysis, producing fructose-1,6-bisphosphate from fructose-6-phosphate. |
|  | *ADH1-ADH5* | *ADH2-ADH4*: early<br>*ADH5*: late | Alcohol dehydrogenase isoenzymes. Dominant fermentative alcohol is *ADH1* (136). Responsible for reoxidation of NADH to NAD<sup>+</sup> which is a required cofactor in the metabolism of glyceraldehyde-3-phosphate in glycolysis. *ADH2* is a non-dominant isoenzyme of alcohol dehydrogenase. Typically repressed by glucose and converts ethanol to acetaldehyde (136). Overexpressed in some wine strains (137). |

*Genes expressed higher in early fermentation in both the 2017 and 2019 vintages.*
| Wine process | Gene | Gene product function |
| --- | --- | --- |
| Glycolysis and fermentation | *HXT1* | Low affinity hexose transporter. |
|  | *HXK2* | Hexokinase that phosphorylates glucose in the first irreversible step leading to glycolysis. |
|  | *PFK1, PFK2* | Phosphofructokinases that catalyze the first irreversible reaction specific to glycolysis, producing fructose-1,6-bisphosphate from fructose-6-phosphate |
| Acetate metabolism | *ALD4-6* | Aldehyde dehydrogenase isoenzymes that produce acetate as a byproduct when acetaldehyde is metabolized. *ALD6* encodes the main isoenzyme responsible for acetate production in wine (96). Aldehyde dehydrogenase isoenzymes *ALD4* and *ALD5* are expressed when ethanol is the carbon source and are not typically associated with wine fermentation. Acetate contributes the majority of volatile acidity associated with negative organoleptic properties in wine (138). |
|  | *PDR12* | Plasma membrane ABC transporter. Required for development of resistance to weak organic acids, including acetate (139). |
| Cell cycle | *BAT1* | Expressed in logarithmic phase (140). |
|  | *CWP1* | Expressed in the S/G2 phase of cell cycle (140). |
| Nitrogen metabolism | *GNP1* | High-affinity glutamine permease that also transports leucine, serine, threonine, cysteine, methionine and asparagine. |
|  | *MUP1* | High-affinity methionine permease that is also involved in cysteine transport. |
|  | *CAR1, CAR2* | Involved in arginine catabolism. Arginine is the most abundant amino acid in grape must after proline (66) and is used in protein synthesis during fermentation (141). |
|  | *YPQ1* | Vacuolar transporter for arginine and lysine. Unused arginine is stored in the vacuole for later use (141). |
| Ehrlich pathway | *BAT1, ARO8* | Catalyzes transamination of amino acids, the product of which cannot be redirected to central carbon metabolism and so is excreted as fusel acid or fusel alcohol (142). Overexpression of *BAT1* increases the concentration of isoamyl alcohol, its acetate ester, and isobutanol in wine (142). |
|  | *PDC1* | Catalyzes alpha keto decarboxylation. |
| Glycerol biosynthesis | *GPP1* | Cleaves phosphate from glycerol-3-phosphate. |

*Genes expressed higher in late fermentation in both the 2017 and 2019 vintages.*
| Wine process | Gene | Gene product function |
| --- | --- | --- |
| Nitrogen limitation | *GAT1, DAL80* | Transcriptional activator (*GAT1*) and repressor (*DAL80*) of genes under nitrogen catabolite repression. Expression is inversely correlated, and the detection of both genes as induced in late fermentation likely indicates tight transcriptional regulation of nitrogen metabolism. |
|  | *DAL2-DAL5, DAL80, DAL82* | Catalyze allantoin degradation and expression is under nitrogen catabolite repression. |
|  | *MEP2* | Ammonia permease and expression is under nitrogen catabolite repression. |
|  | *GAP1* | Amino acid permease. |
|  | *PTR2* | Peptide permease. |
|  | *AVT3, AVT4* | Vacuolar amino acid exporters that mobilize internal nitrogen stores for cell maintenance during stationary phase. Expression is under nitrogen catabolite repression. |
| ubiquitin-mediated selective protein degradation | *RPN4* | Transcription factor that induces expression of proteasome genes. |
|  | *TMC1* | Effector of proteotoxic stress induced by nitrogen limitation, weak acid, and misfolded proteins. Target of *RPN4*. |
|  | *UBC8, VID24* | Negative regulators of fructose-1,6-bisphosphate through ubiquitination (*UBC8*) and vacuolar targeting (*VID24*). |
|  | *UBC1, UBC5, UBC7, UBC13* | Ubiquitin conjugating enzymes. |
|  | *UBI4, CUZ1* | Involved in the ubiquitin-proteasome pathway. |
| Autophagy | *ATG2, ATG4, ATG7-ATG12, ATG14, ATG32, ATG40* | Proteins involved in autophagy. Autophagy is a key response to nutritional limitation that allows cells to maintain homeostasis (143). Nitrogen starvation leads to the largest autophagic response in yeast. |
| Erlich pathway | *GRE2* | Final step of pathway where fusel aldehydes are oxidized or reduced into fusel acids or alcohols (142). |
| Carbon limitation | *SNF3* | Plasma membrane low glucose sensor involved in regulating glucose transport |
|  | *SKS1* | Serine/threonine kinase involved in the adaptation to low glucose via *SNF3*-independent signaling |
|  | *PGM2* | Phosphoglucosmutase. Catalyzes a key step in hexose metabolism. Induced in response to glucose limitation and ethanol stress (144). |
|  | *HXK1* | Hexokinase that phosphorylates glucose or fructose in the first irreversible step leading to glycolysis. Under glucose-induced repression. |
|  | *HXT4, HXT6* | Hexose transporters required at the end of alcoholic fermentation. |
| Trehalose and glycogen | *TSL1, NTH1, ATH1* | Involved in trehalose synthesis (*TSL1*) and degradation (*NTH1, ATH1*). Trehalose acts as a storage carbohydrate for cell maintenance in non-growth conditions (145, 146), bolsters membrane integrity by displacing ethanol (146), and protects proteins from denaturation (147). Trehalose recycling is an important component of stress response (148). |
|  | *GLG1, GSY1, GSY2, IGD1, GPH1, SGA1, GAC1, GIP1, YPI1, PIG2, GLC8* | Involved in glycogen accumulation (*GLG1, GSY1, GSY2, IGD1*), degradation (*GPH1, SGA1*), and metabolism (*GAC1, GIP1, YPI1, PIG2, GLC8*). Glycogen accumulates during nutrient abundance and is metabolized during stationary phase and nutrient deprivation (145). Glycogen recycling is an important component of stress response (148). |
| Cell wall integrity | *PIR3, SED1, SLT2* | Target genes of the cell wall response to ethanol. *PIR3* is required for cell wall stability and induced in part by *SLT2*. *SED1* is a stress-induced cell wall structural protein (*SED1*) (149). |
|  | *PKH2, YPS1, PST1, KRE1* | General cell wall integrity response. |

Beyond these previously defined core gene expression patterns, gene expression signatures indicative of less understood processes within these fermentations are also observed, which may be linked to the industry-like environment these studies were performed in. These patterns of gene expression included signatures of nutrient limitation in early fermentation and polyol metabolism in late fermentation that were both consistent with interactions with non-*Saccharomyces* organisms. We also find signatures of concurrent hypoxic and anoxic metabolism that suggests differential availability of oxygen for some yeast populations throughout fermentation. In association, we observe mounting gene expression that is likely involved in mitigating oxidative and other stresses. Finally, few vintage-specific differences can be found, but those we do identify highlight gene expression patterns that could be linked to altered fermentation outcomes. We discuss these observations below.

### Nutrient limitation in early fermentation

While gene expression data supports logarithmic growth at 16 hours post-inoculation (**Table 1, Figures S1-S2**), at this early timepoint there is also evidence for the expression of genes that are typically up regulated in response to nutrient limitation. *PHO5* and *PHO89* are phosphate transporters that are induced during phosphate starvation (52), both of which are expressed in early fermentation, along with *PHO90*. Phosphate limitation can cause stuck fermentations, as phosphate is critical for cellular function as a component of ATP, nucleotides, sugars, lipids, and macromolecules such as proteins (49, 53). Given that all these Pinot noir fermentations went to completion, and that the majority of glucose was converted to ethanol after the 16hrs time point, induction of genes encoding phosphate transporters early in fermentation is not likely associated with phosphate starvation. Instead, it may be a response to the presence of non-Saccharomyces yeast, as co-cultivation of *S. cerevisiae* with *Torulaspora delbrueckii* led to the induction of a gene encoding a high-affinity phosphate transporter (PHO84) after only three hours of fermentation (40). Enological co-culture of S. cerevisiae with organisms such as *Hanseniaspora guilliermondii* and *Brettanomyces* has also been linked to induction of genes involved in vitamin biosynthesis in fermentation (54, 55), which could be indicative of increased nutrient competition and depletion of some nutrients early in fermentation. We similarly observe induction of genes that encode enzymes involved in biosynthesis of B vitamins in early fermentation, including BIO2 (biotin biosynthesis), RIB3 and RIB4 (riboflavin biosynthesis), PAN6 (pantothenate synthesis), SPE3 and SPE4 (pantothenic acid synthesis), and MIS1 and FOL1 (folate biosynthesis). In addition, THI21 was induced, which is involved in thiamine biosynthesis. As with phosphate, this may be related to the presence of metabolically active non-*Saccharomyces* microorganisms that have been detected in all of these fermentations (56). We expect that continued work using industry-like fermentations across grape varieties and yeast strains, as well as controlled fermentations using reconstituted microbial consortiums, will be critical for understanding the relevance of these gene expression signatures to wine fermentation outcomes. If understood, such interactions could potentially be addressed through timely nutrient additions to a fermentation to achieve desired outcomes.

### Evidence of varied gene expression patterns linked to oxygen exposure during fermentation

A wine fermentation is generally regarded as an anaerobic process given that the carbon dioxide (CO*_2_*) produced as a byproduct of ethanol fermentation protects must from dissolved oxygen (57). Yet, within anaerobia, there is an important distinction between hypoxic (low oxygen) and anoxic (no oxygen) conditions. In a fermentation, it is expected that molecular oxygen (O*_2_*) is introduced into the grape must by handling processes, including pump overs, that may introduce small amounts of dissolved oxygen into industrial-scale tanks (58). Stratification within a fermentation may also expose local cell populations to different oxygen environments leading to yeast cell populations undergoing different anaerobic processes. Within our data, we found gene expression patterns consistent with different populations of cells experiencing varied levels of oxygen exposure during fermentation. For example, the yeast cell wall undergoes remodeling in response to oxygen availability, which is accomplished in part by regulated expression of cell wall mannoproteins encoded by *CWP1*/*CWP2, DAN1,* and *TIR1*-*TIR4* (59). Specifically, expression of *DAN1* and *TIR1*-*TIR4* occurs reciprocally to expression of *CWP1* and *CWP2*, with the *CWP* genes being expressed in aerobic conditions and *DAN1*/*TIR1*-*TIR4* in anaerobic conditions (59). *DAN1* expression is known to be repressed in aerobic conditions by four independent regulatory mechanisms (60). Interestingly, expression of both *CWP1* and *DAN1*/*TIR1*-*TIR4* was observed in early fermentation samples. Similarly, in early fermentations both *HYP2* and *ANB1* were expressed. These paralogous genes encode translation elongation factor eIF5A and are part of a family of paired genes for which oxygen induces the aerobic isoform and represses the hypoxic isoform (61). *HYP2* is expressed under aerobic growth, while *ANB1* is expressed under hypoxic growth and is tightly regulated by the presence of oxygen (62). Together, these gene expression patterns indicate varied gene expression programs within yeasts that may be explained by differing levels of oxygen exposure.

Among late expressed genes, oxygen-regulated paired isoforms were also expressed, including *COX5A* and *COX5B* that encode a subunit of cytochrome *c* oxidase. Modulated expression of these two isoforms allows *S. cerevisiae* to produce holoenzymes with different catalytic properties in response to oxygen (63). *COX5A* expression declines between 5-1 µmol/L O*_2_*, and is undetectable below 0.25 µmol/L O*_2_* while *COX5B* is undetectable until 0.25 µmol/L O*_2_* (61). Simultaneous induction of both transcripts at the end of fermentation is again consistent with cells experiencing varied levels of dissolved oxygen in fermentation (58). In contrast, of the oxygen-regulated isoform pair *CYC1* and *CYC7* (61), only expression of the hypoxic isoform *CYC7* was detected at the end of fermentation. The break point between expression of isoforms occurs at a higher concentration of 0.5 µmol/L O*_2_* for *CYC1*/*CYC7* than *COX5A*/*COX5B* (61), which may indicate that dissolved oxygen levels did not exceed 0.5 µmol/L and therefore was not permissive to expression of *CYC1*.

In late fermentation, induction of pathways like glycerol degradation and proline metabolism that require oxygen were also observed. Glycerol is a compatible solute involved in combating osmotic stress and redox balance and is primarily produced in early fermentation (64). We found induction of *GCY1* which encodes a glycerol catabolic enzyme used in micro aerobic conditions (65), as well as *RSF2*, a transcriptional regulator of genes that encode proteins required for glycerol-based growth. Proline metabolism genes *PUT1*, *PUT2*, and *PUT4* were also expressed at the end of fermentation. Although proline is an abundant amino acid in grape must, it is a non-preferred nitrogen source of yeast and requires oxygen to be metabolized (66). It was further observed that *PUT1* and *PUT2* were induced in a sealed laboratory wine fermentation, but that proline was not metabolized given the absence of oxygen (2). Expression of *PUT1*, *PUT2*, and *PUT4* is regulated by nitrogen catabolite repression (67) and the presence of proline in the absence of other nitrogen sources (68), but is not regulated by the presence of oxygen. Intracellular proline accumulation also protects *S. cerevisiae* from reactive oxygen species associated with ethanol-rich environments (69). While it possible that glycerol and proline were metabolized in late fermentation with oxygen ingress, other processes like nutrient limitation and oxidative stress may also explain the induction of these genes.

Taken together, our gene expression data raises various questions about a distributed gradient of oxygen (hypoxia and anoxia) in the fermentation environment that may induce varied gene expression across the cell population. This could lead to yeast sub-populations undergoing varied metabolic outputs or having different levels of ethanol tolerance due to the role of oxygen in these processes (70, 71). In the future, single cell sequencing technologies combined with continuously monitored dissolved oxygen assays may help resolve these questions. From a production perspective, in industrial fermentations, even those that employ pump over systems and therefore maintain mixing and better homogeneity, there is a gradient of dissolved oxygen in the fermentation tank wherein oxygen concentration is higher toward the top of the vessel (58). This suggests that heterogeneous gene expression profiles in response to oxygen would likely exist in these environments too. This is also an important fact to consider, as oxygen additions during fermentation are known to influence both fermentation and sensory outcomes. For example, in late fermentation, a single oxygen pulse increases the rate of fermentation mediated by ergosterol biosynthesis (70). Similarly, oxygen additions at different stages of fermentation differentially impact wine aroma compound formation like volatile thiols and esters; however, this appears to occur in a strain-dependent manner (71). This knowledge, combined with the impact of oxygen addition on fermentation outcomes, raises the idea that timely addition of oxygen may be a way to control fermentations rates and formation of wine aromas, which would be a tool easily accessible to winemakers.

### Mitochondria and fermentation

In late fermentation, our gene expression analyses find a striking enrichment of pathways involved in mitochondrial biogenesis and function, as well as oxidative phosphorylation (**Figure 2C, Figure S3-S4**). Substantial metabolic investment in mitochondrial systems suggests a critical role for mitochondria late in fermentation. However, what that role is remains unclear, as limited research has been conducted on the mitochondria during enological fermentation, likely because of both low oxygen conditions and the Crabtree effect in fermentative metabolism (72, 73). While some studies that profile the transcriptome of primary fermentation either find no evidence for, or make no comment on enrichment for oxidative metabolism at the end of fermentation, many studies have found induction of mitochondrial genes, particularly those encoding oxidative phosphorylation. This includes fermentations conducted under nitrogen limitation (6), lipid limitation (74), and standard laboratory conditions (3). Interestingly, under lipid limitation, oxidative phosphorylation was induced in the exponential phase of growth as opposed to the end of fermentation (74). Given the role of membrane lipid composition in combating ethanol-induced membrane permeability (75), and the accumulation of reactive oxygen species during ethanol exposure (76), induction of the respiratory chain may mitigate reactive oxygen species that are abundant at the end of fermentation. Nonetheless, the recurrence of these gene expression patterns in our studies and previous laboratory experiments suggest that cells are investing in mitochondrial systems during fermentation.

One potential reason for late induction of mitochondrial systems is that glucose-limitation relieves the Crabtree effect. This may lead to induction of oxidative phosphorylation genes that change metabolism in a nutrient-limited environment to one that generates the largest amount of ATP per unit of glucose (77). In this way, an investment in mitochondrial infrastructure during late fermentation may be a starvation adaptation in which *S. cerevisiae* uses oxidative phosphorylation to harness the largest fraction of energy possible from remaining carbohydrate sources. However, this strategy is predicated on availability of molecular oxygen, which is required for the induction and function of the respiratory apparatus (78, 79). A second reason for mitochondrial gene expression may be related to the fact that meiosis and sporulation related genes were enriched at the end of fermentation (**Figure 2C**, **Figure S3-S4**). Induction of meiosis likely occurs to produce spores resistant to the challenges of nutrient limitation and stress (80). Interestingly, mitochondrial biomass is a predictor for meiosis (81), and components of the respiratory chain are required for initiation of sporulation (82), providing another potential process that may underlie mitochondrial investment in late fermentation. Related to this fact, a propensity for yeast to undergo meiosis at this stage of vinification underlies fast adaptive genomic evolution of *S. cerevisiae* (83), suggesting this may be an important acquired trait that allows yeast to successfully survive the wine environment.

Mitochondria also fulfill other critical roles in fermentation unrelated to respiration. For example, mitochondria play a role in sterol uptake and transport under strictly anaerobic conditions (84), and mitochondria quench reactive oxygen species especially during ethanol stress (85). While we did not observe induction of specific genes related to sterol biology and found induction of different genes related to reactive oxygen species than those previously identified (see next section), these processes may also be linked to increased mitochondrial gene expression. Regardless of the role played by mitochondria in late fermentation, the striking and consistent induction of these genes in fermentations signals that more research is needed to understand the role of mitochondria in fermentation.

### Thioredoxins and glutathione system activity throughout fermentation

The reducing environment of the cytosol in *S. cerevisiae* is key to various cellular functions, including deoxyribonucleoside triphosphate synthesis and the elimination of toxic compounds, including oxidants generated through cellular metabolism (86, 87). Key to maintaining redox balance are the thioredoxin (TRX) and glutathione (GSH) thiol-reductase systems. For example, proper redox homeostasis is required to maintain the redox status of cysteine residues, which are essential for the function of numerous enzymes, protein receptors, and transcription factors. Similarly, redox homeostasis within cells aids to balance pools of reduced and oxidized pyridine nucleotide cofactors (NAD+/H, NADP+/H) that are essential to numerous metabolic reactions. Reactive oxygen species (ROS) can alter this redox balance causing oxidative stress and direct or indirect ROS-mediated damage of nucleic acids, proteins, and lipids. While typically associated with respiratory metabolism, ROS can be generated throughout fermentation, in particular by superoxide anions and peroxides (76, 88, 89). ROS may also be created by acetaldehyde, an intermediate in ethanol production (90).

In early fermentation, we see induction of genes involved in the thioredoxin system, such as *TRX1* and *TRR1*. Expressed targets of *TRX1* included *RNR1-RNR4* (91), genes encoding ribonucleotide-diphosphate reductase required for DNA synthesis and cell cycle progression, as well as *MET16*, which encodes an enzyme required for sulfate assimilation (92). We further observed genes encoding Trx1 target peroxidases (*TSA1*) and peroxiredoxins (*AHP1*) constitutively expressed throughout fermentation along with superoxide dismutases (*SOD1*, *SOD2*). An additional source of ROS are peroxisomes, which may generate hydrogen peroxide in early fermentation via beta-oxidation of fatty acids. *CTA1*, which encodes a peroxisomal catalase, and *ANT1*, which encodes a peroxisomal transporter involved in beta-oxidation of fatty acids, were expressed in early fermentation. A major factor used to maintain redox balance is NADPH, which provides reducing potential for the thioredoxin system. It has been shown that metabolic intermediates in glycolysis can be re-routed to the pentose phosphate pathway to generate NADPH in response to oxidative stress (93–95). We found that the pentose phosphate pathway was enriched among genes expressed in early fermentation (**Figure 2C**, **Figure S1**-**S2**), which includes *GND1*, an enzyme that catalyzes NADPH regeneration and is required for the oxidative stress response. Other expressed genes that encode enzymes acting downstream of *GND1* in the pentose phosphate pathway included *RPE1*, *TLK1*, *TLK2*, and *TAL1*.

Central to the glutathione (GSH) thiol-reductase system is glutathione, an abundant tripeptide conserved throughout eukaryotic and prokaryotic cells with a critical role in redox control, but its physiological role is both diverse and debated (95). We observed that genes encoding enzymes involved in the degradation (*DUG1* and *DUG2*), import (*OPT1*), and biosynthesis (*GSH1* and *GSH2* in the 2017 vintage) of glutathione were expressed in early fermentation. Additional generation of NADPH in early fermentation may be supported by the transformation of isocitrate to alpha ketoglutarate via *IDP1* in the mitochondria, and export via *YMH2*, as both genes were also expressed. Genes encoding aldehyde dehydrogenases *ALD5* and *ALD6* are similarly expressed in early fermentation, both of which may regenerate NADPH through the transformation of acetaldehyde to acetate. *ALD6* is the dominant isoenzyme responsible for acetate production in wine (96).

We further observed induction of genes involved in glutathione-mediated ROS mitigation in late fermentation. For example, a gene encoding cytosolic glutaredoxin (*GRX1*) was expressed in late fermentation. Unlike glutaredoxins in other species (e.g., mammals), yeast glutaredoxins do not function as deglutathionylase enzymes (97). Instead, induction of *GRX1* increases resistance to hydroperoxides by catalytically reducing hydroperoxides through glutathione conjugation and using the reducing power of NADPH (98). In addition, the cytosolic peroxidase *GPX1* was expressed. *GPX1* uses both glutathione and thioredoxin, in combination with NADPH, for reducing power (99). *GPX1* is known to be expressed by glucose and nitrogen starvation (100), which coincides with peak peroxide formation in yeast during wine fermentation (88). While our gene expression data support a role for cytoplasmic glutathione during late fermentation, genes encoding mitochondrial peroxidin (*PRX1*) and thioredoxin (*TRX3*) were also expressed. Prx1 buffers the mitochondria from oxidative stress and is reductively protected by glutathione, thioredoxin reductase (Trr2), and Trx3 (101). Taken together, these results suggest that cytoplasmic and mitochondrial systems may be integral to combating increased oxidative stress at the end of fermentation.

Glutathione is also important for maintenance of cellular function via other systems. For example, methylglyoxal is a byproduct of glycolysis, a reduced derivative of pyruvic acid, that may account for up to 0.3% of glycolytic carbon flux in *S. cerevisiae* (102). We found that *GLO2*, an enzyme that catalyzes methylglyoxal degradation in a glutathione dependent manner, was expressed in late fermentation, as were glutathione-independent systems involved in the degradation of methylglyoxal (*GRE2*/*GRE3*). Genes that encode proteins involved in glutathione homeostasis were also expressed at the end of fermentation, including *GEX1* that encodes a proton:glutathione antiporter (103, 104). *GEX1* is known to be induced during oxidative stress (103) and modulates formation of the aromatic thiol 3-mercaptohexan-1-ol from its glutathionylated precursor in wines such as Sauvignon Blanc (104). Conversely, an induction of a gene that encodes an enzyme that cleaves glutathione (*GCG1*) was observed and may be involved in apoptotic signaling via ROS accumulation (105).

Together, these gene expression patterns highlight how intertwined redox homeostasis is with almost all core metabolic processes in *S. cerevisiae*, as most pathways require oxidation or reduction by a pyridine nucleotide cofactor during at least one reaction. For example, NAD+/H and NADP+/H participate in 740 and 887 biochemical reactions through interactions with 433 and 462 enzymes, respectively (106). It is also well documented that experimental perturbation of both NAD+/H and NADP+/H leads to changes in aroma compounds in wine and other fermented beverages (107–110). The observations presented herein conserved across many Pinot noir fermentations involving genes engaged in redox balance and mitigation of oxidative stress via thiol-reductase systems offers further evidence for the importance of these systems. These findings provide motivation for future studies of these systems in the context of wine production, which would include control measures to aid cellular control of redox and mitigate oxidative cellular stress.

### Stress-associated gene expression during fermentation

During fermentation, *S. cerevisiae* has to adapt to a continually changing stress landscape. Macro- and micronutrients become limiting as ethanol concentrations increase and, as discussed above, production of acetaldehyde and other metabolic processes generate oxidative stress. To accommodate this dynamic environment, *S. cerevisiae* wine strains express genes that overlap with, but are distinct from, the stress response of laboratory strains. (7, 111). In accordance with previous studies (2, 3), we found partial overlap between genes expressed in fermentation and those involved in the environmental stress response (ESR) in laboratory strains. Specifically, 16 ESR genes were expressed at the beginning of fermentation and 78 ESR genes at the end of fermentation. This matches observations in synthetic must where stress genes were induced upon entry into stationary phase (2). Stress-related genes expressed at the beginning of fermentation were enriched for Gene Ontology pathways involving carbohydrate transmembrane transport (mannose, fructose, glucose, hexose) and NADP regeneration **(Figure 3A)**, while stress-related genes expressed at the end of fermentation were enriched for oxidation-reduction process, generation of precursor metabolites and energy, energy reserve metabolic process, and glycogen metabolic process **(Figure 3B).**

**Figure 3.**
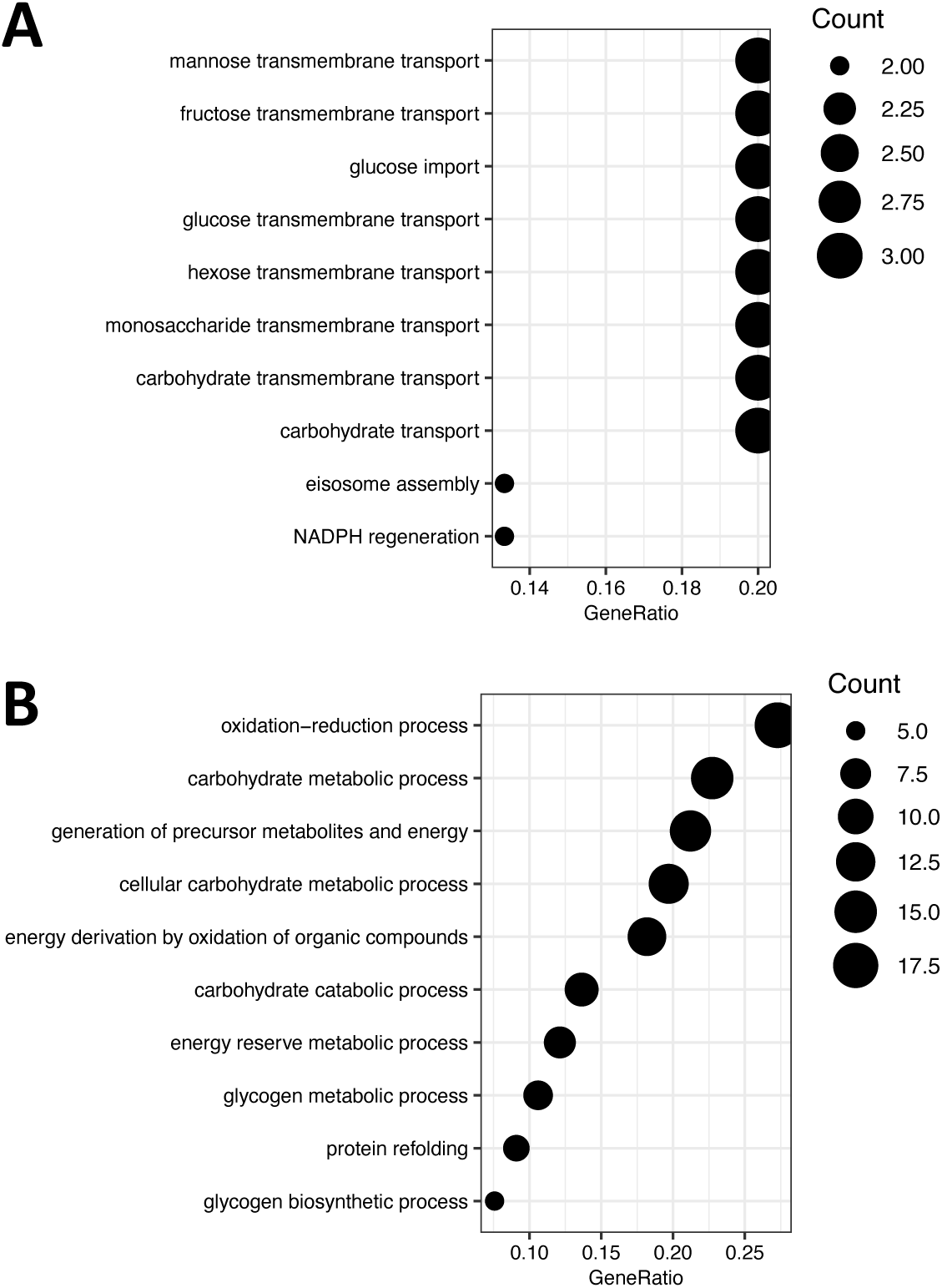
Pathways enriched among genes differentially expressed across fermentation that are shared with the environmental stress response (ESR). **A.)** Of 16 genes that overlap with the ESR and were expressed in early fermentation, pathways related to carbohydrate metabolism were enriched. **B.)** Of 78 genes that overlap with the ESR and were expressed in later fermentation, pathways related to oxidation-reduction and carbohydrate metabolism were enriched.

A recent study investigated fermentation of Riesling grape must at laboratory scale without addition of oxygen (3). Using microarray analysis at five time points in fermentation, the authors defined a fermentation stress response (FSR) as those genes that are induced at any point in fermentation and do not return to baseline (3). The FSR is differentiated from the ESR and common stress response because adaptation is not observed over time through gene expression returning to pre-stress transcription levels (3, 7, 111). Of the 223 genes induced in the FSR, 84 were observed to be expressed in mid- or late-fermentation. Of these 84 genes, 43 overlap with genes expressed in other stress responses as defined in (3), including 16 with the ESR and 14 with the common stress response. Of the 41 genes that overlap with the FSR, many were related to the challenging nutrient environment in wine, including glucose limitation (*NRG1*, *SKS1*, *HXT6*, *VID24*), nitrogen limitation (*MEP2*, *GAP1*, *PTR2*, *AVT4*, *VBA2*), vitamin limitation (*MCH5*, *VHR1*), and stress caused by heat, salt, protein mis-folding, and cell wall defects (*GAC1*, *RPI1*, *JID1*, *PSR2*). This suggests that multiple stress pathways are simultaneously activated by the challenging environment that *S. cerevisiae* encounters in wine fermentation, which likely defines the described fermentation stress response. Many genes identified in the FSR, and expressed in this study during fermentation, remain uncharacterized (*YPR152C*, *YBR085C-A*, *YDL024C*, *YDR042C*, *YMR244W*, *YLL056C*), offering gene targets of future investigation related to adaptation to the fermentation and wine environment.

### Polyol metabolism in late fermentation

Polyols, also called sugar alcohols, have recently been shown to be produced by non-*Saccharomyces* yeasts and by fructophilic lactic acid bacteria such as *Lactobacillus kunkeei* during fermentation (112, 113). Combined with other spoilage organism associated metabolites, these compounds can have a significant impact on wine quality (114). Mannitol is one such polyol and a non-preferred sugar that can be metabolized by *S. cerevisiae* (115–117). In *S. cerevisiae*, transporters encoded by *HXT13* and *HXT15-17* were found to facilitate mannitol and sorbitol transport (116). In our data, we observed induction of the mannitol transporter *HXT13* in both vintages, along with the mannitol dehydrogenase *MAN2*, which together indicate that mannitol may be present and metabolized by *S. cerevisiae* at the end of fermentation (**Figure 4**). In line with this, although eukaryotic transcriptional profiling via 3′ Tag-seq was performed (see methods), *L. kunkeei* transcripts were detected in some fermentations, which is one potential source of mannitol production. These data raises the possibility of mannitol consumption by *S. cerevisiae*, demonstrating metabolic flexibility for carbon sources late in fermentation.

**Figure 4.**
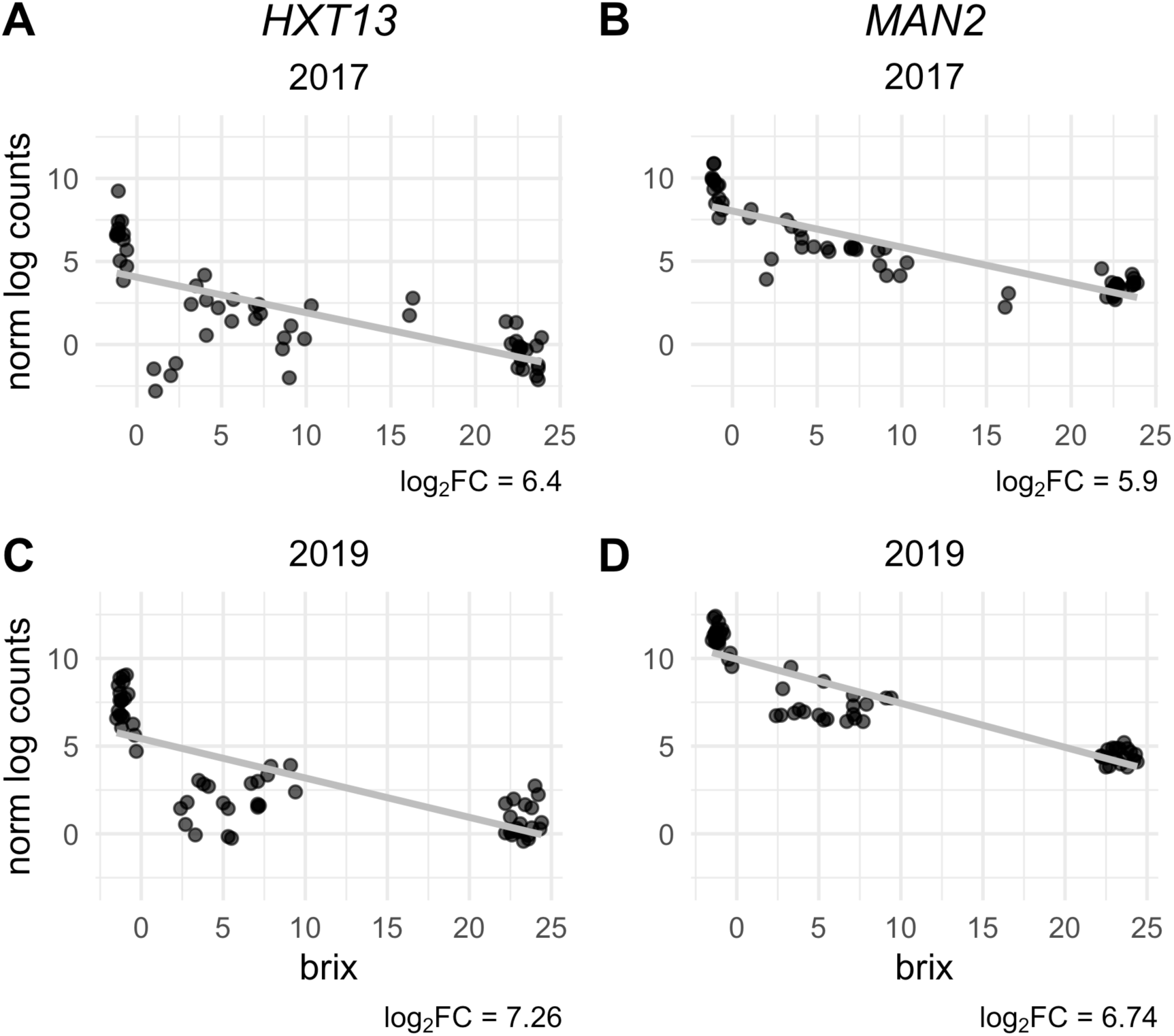
Normalized log gene expression counts for genes involved in mannitol transport and degradation. HXT13 and MAN2 expression is graphed in the 2017 (**A, B**) and 2019 (**C, D**) vintages. MAN2 and HXT13 were the top most expressed genes at the end of fermentation in 2019, and only fell behind HXT4 in the 2017 vintage. Grey lines indicate a linear model fit to normalized counts.

Notably, *L. kunkeei* can influence *S. cerevisiae* metabolism beyond expression of genes for non-preferred carbon sources. Via production of acetic acid, and possibly other compounds, *L. kunkeei* has been shown to induce the [GAR+] prion phenotype in *S. cerevisiae* thereby shifting carbon metabolism away from hexoses (118, 119). Given that the presence of *L. kunkeei* RNA was detected in the 2017 vintage, we tested for the presence of the [GAR+] phenotype in the 2019 vintage via cell culture (119). We did not detect the presence of the [GAR+] prion in any fermentation tested in the 2019 vintage. While the absence of the [GAR+] phenotype in the 2019 vintage does not preclude its presence in the 2017 vintage, consistent gene expression for mannitol transport and degradation in both vintages suggests that *S. cerevisiae* may be metabolizing mannitol in these Pinot noir fermentations due to the presence of *non-Saccharomyces* organisms, including *L. kunkeei*.

### Vintage-specific differences

From our data analyses, there are 717 genes and 375 genes differentially expressed in the 2017 and 2019 vintages, respectively. The majority of these genes were members of pathways enriched among all fermentations (**Figure 2C**, **Figure S1**-**Figure S4**). Using Gene Ontology enrichment analysis, no molecular function, cellular compartment, or biological process was enriched in either vintage that was not also enriched in both vintages. This suggests these differences may be largely due to sequencing depth or variations in the gene expression within these pathways and not differences in the overall biology of *S. cerevisiae*. Still, signatures indicative of vintage-specific effects were observed, some of which may impact the sensory attributes of wine. For example, glycerol is an important fermentation byproduct that can contribute to the mouth feel of wine (120). *S. cerevisiae* uses glycerol biosynthesis to generate NAD+, a required cofactor for glycolysis, when NAD+ levels are not sufficiently replenished through fermentation (121). During glycerol biosynthesis, enzymes encoded by *GPD1* and *GPD2* convert dihydroxyacetone phosphate into glycerol-3-phosphate (122). Both *GPD1* and *GPD2* were expressed in early fermentation in the 2017 vintage, but not in the 2019 vintage. A second example involves genes encoding the fluoride transporters Fex1 and Fex2, which were expressed in late fermentation across all fermentations in the 2019 vintage. Fluoride is a toxic anion that *S. cerevisiae* exports via two plasma membrane transporters to avoid cell damage (123), which in excess can cause slow or stuck fermentation (124). Although fluoride is ubiquitous in terrestrial and aquatic environments (123), application of the insecticide Cryolite, which contains fluoride, has caused problematic fermentations in California vineyards (124). Currently, the reasons for these gene vintage-specific expression patterns are not known.

Finally, it was observed that genes of currently unknown function were differentially expressed in the two vintages assayed. Using a log*_2_* fold change cut off of two, 14 genes in the 2017 vintage and seven genes in the 2019 vintage were of unknown function. Across both vintages, more genes of unknown function were expressed in late fermentation than in early fermentation (10 in 2017, 5 in 2019). Knowledge of the specific pathways expressed in late fermentation due to the stressful, nutrient-limited conditions, offers clues to the potential functions of these genes that could be explored in future work.

## Conclusion

In this study, we present a gene expression analysis across 40 pilot-scale fermentations of California Pinot noir wine using grapes from 10 vineyard sites and two vintages. The fermentations were diverse with different kinetics, initial chemical conditions, and microbial communities (56). Yet among this diversity, we detected a core gene expression program by *S. cerevisiae* that is largely consistent with that observed at laboratory scale (2–4). Given that there are many genes consistently expressed across these Pinot noir fermentations from diverse vineyards, members of this core fermentation gene program represent strong candidates for future study to impact wine outcomes, e.g. through manipulating redox balance (107–110). Excitingly, this includes a large number of genes with unknown function that through investigation may provide new insights into the biology of *S. cerevisiae*.

The largest deviations from benchtop fermentations are likely attributed to activities of non-*Saccharomyces* organisms, but more research is needed to understand these complex ecological interactions and their impact on fermentation. The gene expression signatures around oxygen presence and metabolic availability also warrants further research, in particular into the role of the mitochondria in late fermentation (3, 6, 74). While we detected few vintage-specific differences between fermentations, we expect there are vineyard-site specific deviations from the consistent patterns of gene expression described herein. Given the variability in fermentation kinetics with respect to time of sampling, new methods will likely be needed to resynchronize stages of fermentation to enable cross-vineyard comparisons (4). Future work is also needed to extend these observations to other grape varieties and *S. cerevisiae* wine strains, which will define both the shared and unique facets of the core gene expression program in *S. cerevisiae* linked to these variables. With such information, we can address the impact of an industrial wine fermentation environment on *S. cerevisiae* gene expression and define approaches that can be used to manage commercial fermentation outcomes.

## Methods

### Grape preparation and fermentation

The wine making protocol used in this study was described previously (23, 48). The grapes used in this study originated from 10 vineyards in six American Viticulture Areas in California. All grapes were Pinot noir clone 667 rootstock 101-14. We harvested grapes at approximately 24 Brix and transported the fruit to University of California, Davis Pilot Winery for fermentation. We performed separate fermentations for grapes from each site, with two fermentations per site, totaling of 20 fermentations per vintage (40 fermentations total). After harvest, we separated the fruit into half-ton macrobins on harvest day and added Inodose SO*_2_* to 40 ppm. We stored the bins in a 14°C cold room until destemming and dividing of the fruit into temperature jacket-controlled tanks. We performed N*_2_* sparging of the tank headspace prior to fermentation and sealed tanks with a rubber gasket. We cold soaked the must at 7°C for three days and adjusted TSO*_2_* to 40 ppm on the second day. After three days, we increased the must temperature to 21°C and set a programmed pump over timetable to hold the tank at a constant temperature. We reconstituted *S. cerevisiae* RC212 with Superstart Rouge at 20 g/hL and inoculated the must with 25 g/hL of yeast. At approximately 24 hours after inoculation, we adjusted nitrogen content in the fermentations using DAP (target YAN – 35 mg/L – initial YAN)/2, and Nutristart using 25 g/hL. We only adjusted nitrogen if target YAN was below 250 mg/L. Approximately 48 hours after fermentation, we permitted fermentation temperatures to increase to 27°C and added DAP as previously described. Fermentations ran to completion when Brix < 0. We took fermentation samples for Brix measurements and RNA isolation at 16, 64, and 112 hours relative to inoculation. To ensure uniform sampling, we performed a pumpover ten minutes prior to sampling each tank. For RNA samples, we obtained 12mL of juice was obtained and centrifuged at 4000 RPM for 5 minutes. We discarded the supernatant and froze the pellet in liquid nitrogen. We stored samples at −80°C until RNA extraction.

### RNA extraction and sequencing

We thawed frozen yeast pellets on ice, resuspended in 5ml Nanopure water, centrifuged at 2000g for 5min, and aspirated the supernatant. We extracted RNA using the Quick RNA Fungal/Bacterial Miniprep kit including DNAsel column treatment (cat#R2014, Zymo Research). We eluted samples in 30µL of molecular grade water and assessed for concentration and quality via Nanodrop and RNA gel electrophoresis. We adjusted sample concentrations to 200ng/µl and 20 µl sent for sequencing. We used 3′ Tag-seq single-end sequencing (Lexogen QuantSeq) in both the 2017 and 2019 vintage, with the addition of UMI barcodes in 2019. The University of California, Davis DNA Technologies Core performed all library preparation and sequencing.

### Differential expression analysis

We preprocessed samples according to manufacturer recommendations. First, we hard-trimmed the first 12 base pairs from each read and removed Illumina TruSeq adapters and poly A tails. Next, we used STAR to align our reads against *S. cerevisiae* S288C genome (R64, GCF_000146045.2) with parameters --outFilterType BySJout -- outFilterMultimapNmax 20 --alignSJoverhangMin 8 --alignSJDBoverhangMin 1 -- outFilterMismatchNmax 999 --outFilterMismatchNoverLmax 0.6 --alignIntronMin 20 --alignIntronMax 1000000 --alignMatesGapMax 1000000 --outSAMattributes NH HI NM MD --outSAMtype BAM SortedByCoordinate (125). For the 2019 vintage, we used UMI tools to deduplicate alignments (126). We then quantified reads mapping to each open reading frame using htseq count (127). We imported counts into R and filtered to mRNA transcripts. To prepare for differential expression, we used the edgeR function calcNormFactors with default parameters (128). We used limma for differential expression, building a model using Brix values, preparing the data for linear modelling with the voom function, and building a linear model for each gene with lmFit (129). We considered any gene with an adjusted p value < 0.05 as significant. To combat batch effects from different library preparation techniques used in the 2017 and 2019 vintages, we performed differential expression separately on counts from each vintage. We took the union of expressed and repressed genes between vintages, respectively, to generate the final set of differentially expressed genes. We visualized expressed and repressed genes using proteomaps (130), and visualized the intersection of differentially expressed genes between vintages using the R package ComplexUpset (https://github.com/krassowski/complex-upset).

We performed gene set enrichment analysis for genes that were expressed and repressed in both vintages against the Gene Ontology (ont = “ALL”) and Kyoto Encyclopedia of Genes and Genomes (organism = “sce”) databases using the R package clusterProfiler (131).

### Detection of *Lactobacillus kunkeii* in RNA sequencing reads

3′ Tag-seq sequences the tail-end of transcripts that contain poly(A) tails. The majority of transcripts with poly(A) tails are eukaryotic in origin, but given that bacteria perform polyadenylation as a degradation signal (132), a very small subset of transcripts may also originate from bacteria. We identified *Lactobacillus kunkeei* in RNA-seq reads using sourmash gather (133, 134). Using all *L. kunkeei* genomes available in GenBank (08/06/2019), we generated sourmash signatures for each using a k-mer size of 31 and a scaled value of 100. We then used sourmash index to generate a database of *L. kunkeei* genomes, and queried this database using signatures of each RNA-seq sample. To validate findings from sourmash gather, we used BWA mem with default parameters to map a subset of samples against the best matching *L. kunkeei* genome (135).

### Culturing *Saccharomyces cerevisiae* for [GAR+] prion detection

To ascertain whether the [GAR+] prion state was detectable in wine fermentations, we cultured yeast for the prion as performed in (119). We used yeast peptone-based medium containing the designated carbon source, such as YPD (1% yeast extract, 2% peptone, 2% agar, 2% glucose); YPG (1% yeast extract, 2% peptone, 2% agar, 2% glycerol) or GGM (1% yeast extract, 2% peptone, 2% agar, 2% glycerol, 0.05% D-[+] glucosamine hydrochloride). We inoculated yeast from fermentations into each well of a 96 well plate containing 200µl liquid YPD + 34g/ml chloramphenicol, and then grew yeast at 30°C for 48 hours. We then pinned yeast to YPG or GGM plates and grew at 30°C for four days.

### American Viticultural Area (AVA) map construction

We constructed the AVA map featured in **Figure 1** from the UC Davis Library AVA project https://github.com/UCDavisLibrary/ava.

## Data Availability

RNA sequencing data is available in the Sequence Read Archive under accession number PRJNA680606. All analysis code is available at github.com/montpetitlab/Reiter_et_al_2020_GEacrossBrix.

## Acknowledgements

We thank all past and current members of the Runnebaum and Montpetit laboratories for their support of this work, as well as the students and staff of the UC Davis Pilot Winery. T.R. and C.T.B were supported by the Gordon and Betty Moore Foundation’s Data-Driven Discovery Initiative [GBMF4551]. T.R. was supported by the Harry Baccigaluppi Fellowship; Horace O Lanza Scholarship; Louis R Gomberg Fellowship; Margrit Mondavi Fellowship; Haskell F Norman Wine & Food Fellowship; Chaîne des Rôtisseurs Scholarship; Carpenter Memorial Fellowship. The authors would like to recognize financial support from Jackson Family Wines, in addition to support from Lallemand Inc.

---

**Figure S1.**
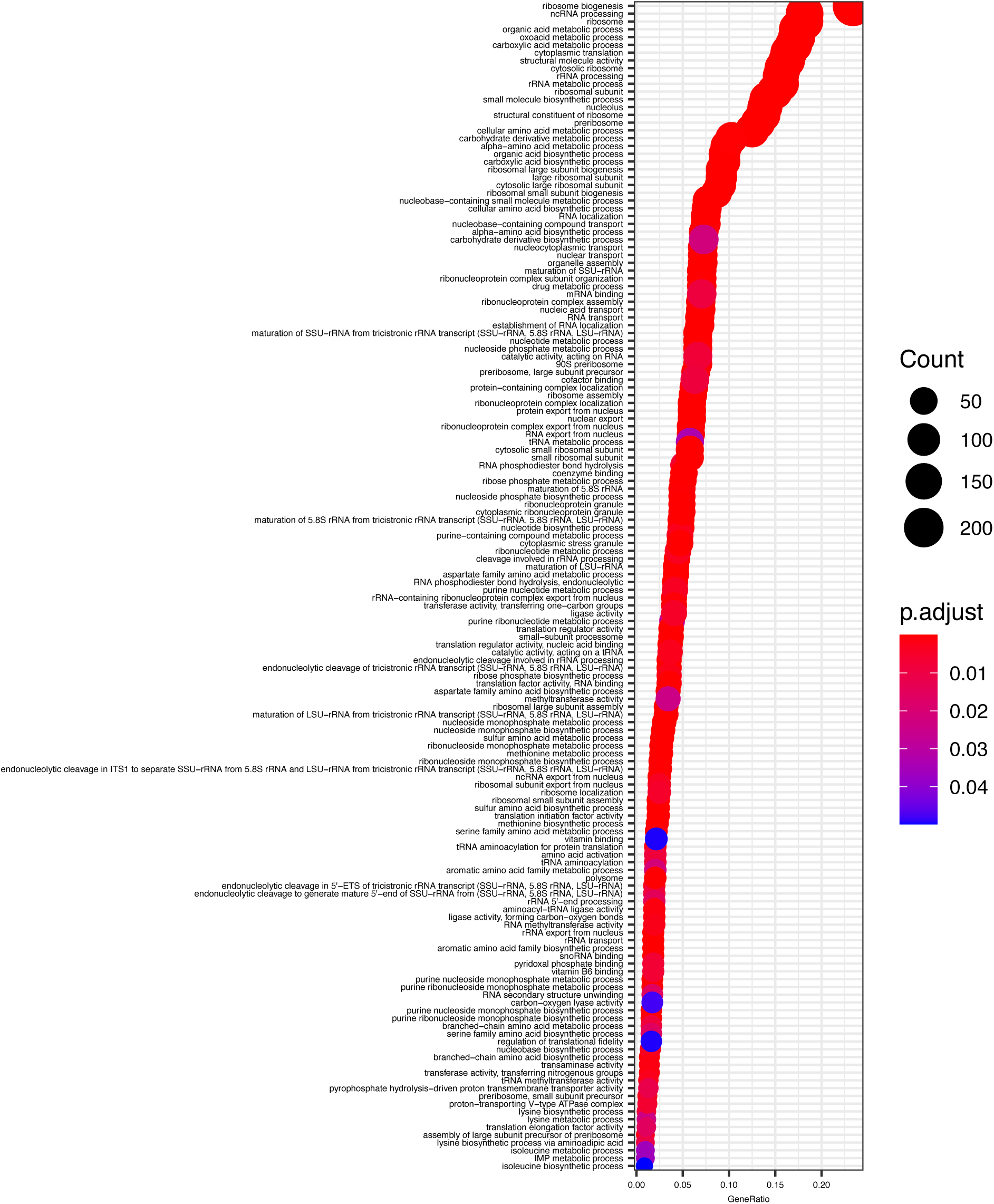
Significantly enriched Gene Ontology (GO) pathways in early fermentation. Pathways from GO categories molecular function, cellular component, and biological process are represented. Significant pathways are defined as p < 0.05 after Bonferroni p value correction.

**Figure S2.**
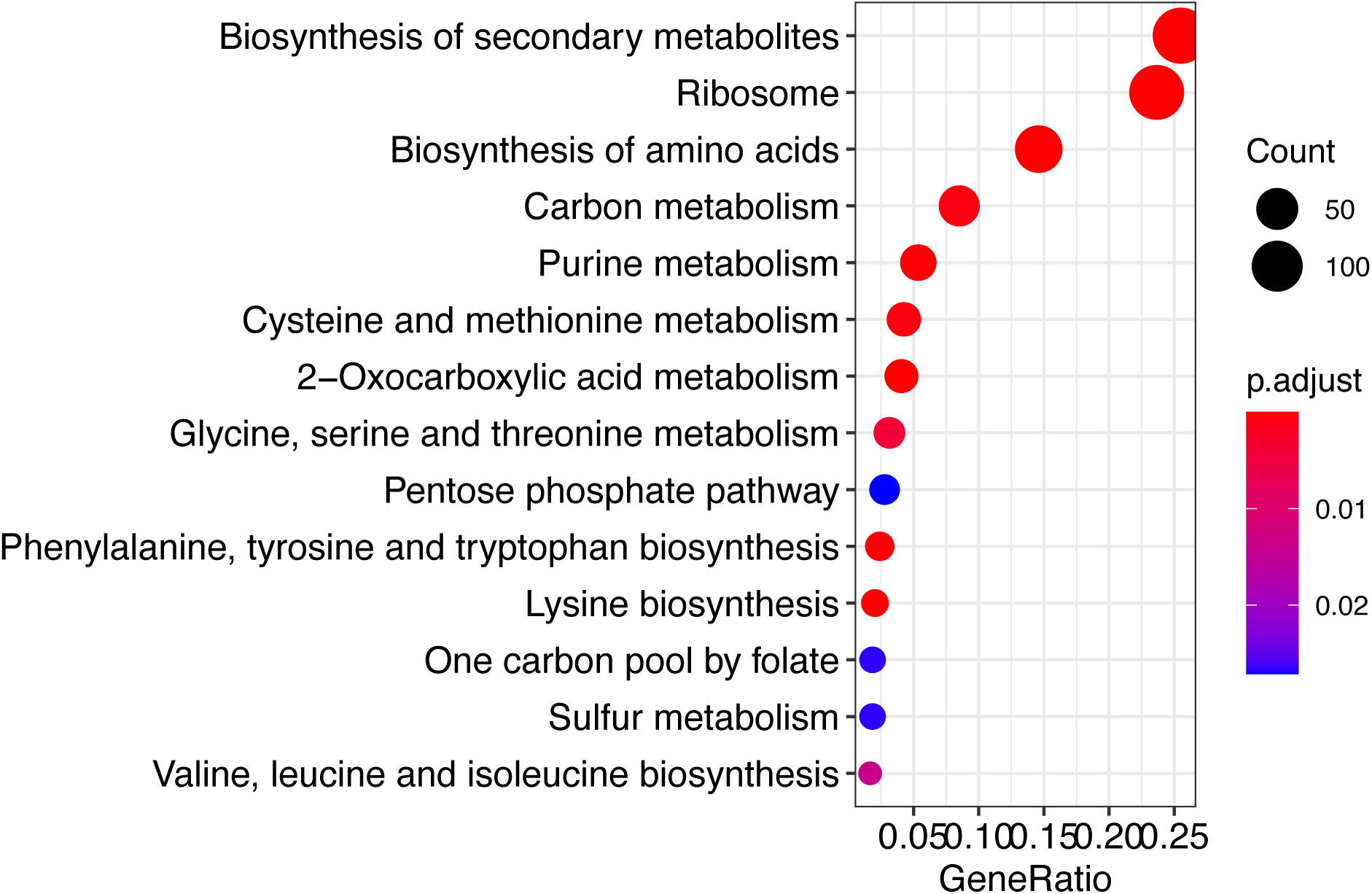
Significantly enriched KEGG pathways in early fermentation. Significant pathways are defined as p < 0.05 after Bonferroni p value correction.

**Figure S3.**
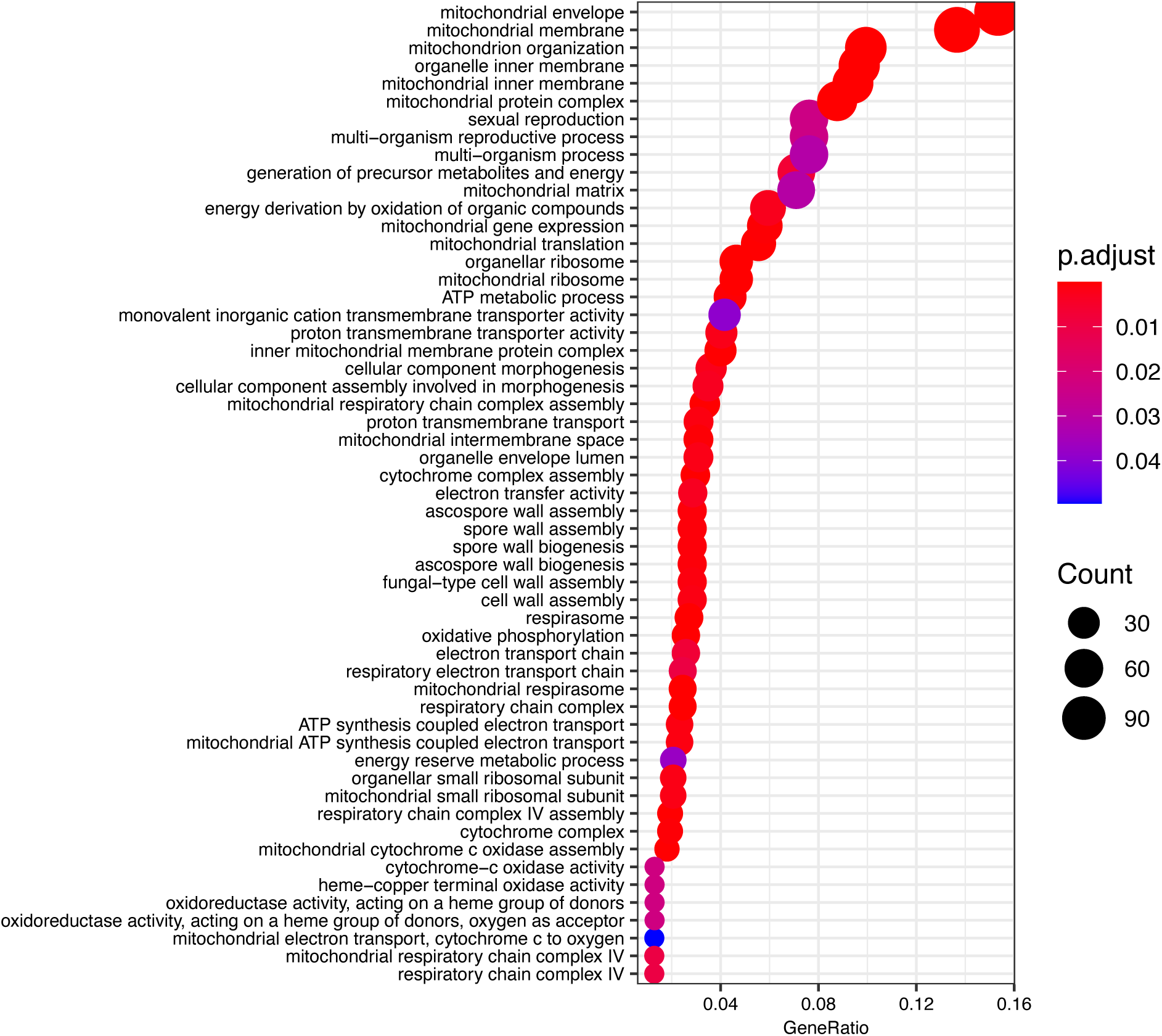
Significantly enriched Gene Ontology (GO) pathways in late fermentation. Pathways from GO categories molecular function, cellular component, and biological process are represented. Significant pathways are defined as p < 0.05 after Bonferroni p value correction.

**Figure S4.**
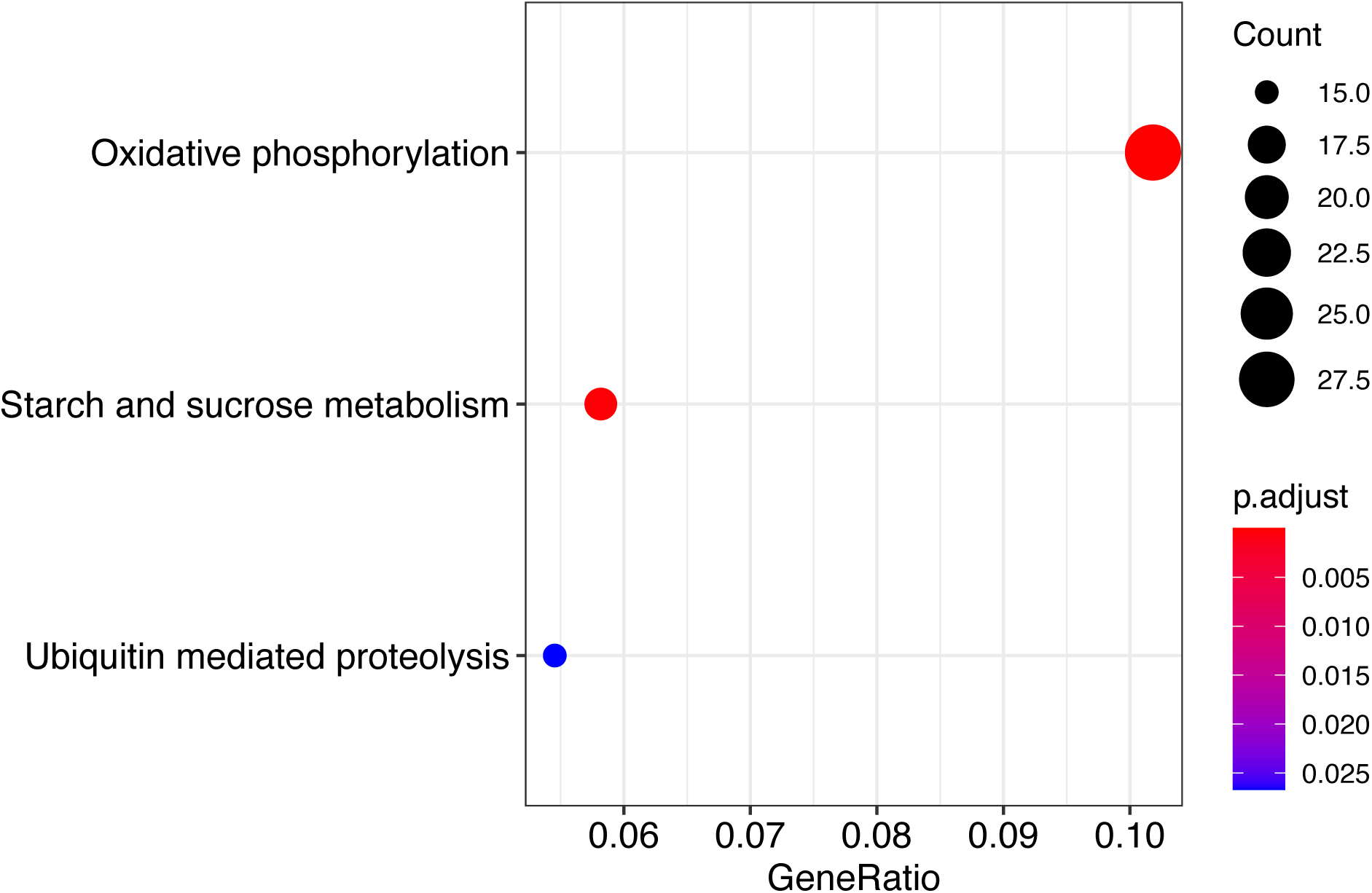
Significantly enriched KEGG pathways in late fermentation. Significant pathways are defined as p < 0.05 after Bonferroni p value correction.

## Notes

https://github.com/montpetitlab/Reiter_et_al_2020_GEacrossBrix

### Competing Interest Statement

The authors have declared no competing interest.

### Summary of Updates

Supplemental files added.

## References

1. Querol A, Fernández-Espinar MT, lí del Olmo M, Barrio E. 2003. Adaptive evolution of wine yeast. International journal of food microbiology 86:3–10.

2. Rossignol T, Dulau L, Julien A, Blondin B. 2003. Genome-wide monitoring of wine yeast gene expression during alcoholic fermentation. Yeast 20:1369–1385.

3. Marks VD, Ho Sui SJ, Erasmus D, Van Der Merwe GK, Brumm J, Wasserman WW, Bryan J, Van Vuuren HJ. 2008. Dynamics of the yeast transcriptome during wine fermentation reveals a novel fermentation stress response. FEMS yeast research 8:35–52.

4. Rossouw D, Jolly N, Jacobson D, Bauer FF. 2012. The effect of scale on gene expression: commercial versus laboratory wine fermentations. Applied microbiology and biotechnology 93:1207–1219.

5. Backhus LE, DeRisi J, Brown PO, Bisson LF. 2001. Functional genomic analysis of a commercial wine strain of Saccharomyces cerevisiae under differing nitrogen conditions. FEMS Yeast Research 1:111–125.

6. Mendes-Ferreira A, Del Olmo M, García-Martínez J, Jiménez-Martí E, Mendes-Faia A, Pérez-Ortín JE, Leao C. 2007. Transcriptional response of Saccharomyces cerevisiae to different nitrogen concentrations during alcoholic fermentation. Applied and Environmental Microbiology 73:3049–3060.

7. Gasch AP, Spellman PT, Kao CM, Carmel-Harel O, Eisen MB, Storz G, Botstein D, Brown PO. 2000. Genomic expression programs in the response of yeast cells to environmental changes. Molecular biology of the cell 11:4241–4257.

8. Puig S, Pérez-Ortín JE. 2000. Stress response and expression patterns in wine fermentations of yeast genes induced at the diauxic shift. Yeast 16:139–148.

9. Bisson LF. 1999. Stuck and sluggish fermentations. American Journal of Enology and Viticulture 50:107–119.

10. Gutiérrez A, Beltran G, Warringer J, Guillamón JM. 2013. Genetic basis of variations in nitrogen source utilization in four wine commercial yeast strains. PLoS One 8:e67166.

11. Patel S, Shibamoto T. 2002. Effect of different strains of Saccharomyces cerevisiae on production of volatiles in Napa Gamay wine and Petite Sirah wine. Journal of agricultural and food chemistry 50:5649–5653.

12. Rossouw D, Næs T, Bauer FF. 2008. Linking gene regulation and the exo-metabolome: a comparative transcriptomics approach to identify genes that impact on the production of volatile aroma compounds in yeast. BMC genomics 9:530.

13. Riou C, Nicaud J-M, Barre P, Gaillardin C. 1997. Stationary-phase gene expression in Saccharomyces cerevisiae during wine fermentation. Yeast 13:903–915.

14. Rossouw D, Bauer FF. 2009. Comparing the transcriptomes of wine yeast strains: toward understanding the interaction between environment and transcriptome during fermentation. Applied microbiology and biotechnology 84:937.

15. Casalta E, Aguera E, Picou C, Rodriguez-Bencomo J-J, Salmon J-M, Sablayrolles J-M. 2010. A comparison of laboratory and pilot-scale fermentations in winemaking conditions. Applied microbiology and biotechnology 87:1665–1673.

16. Walker ME, Nguyen TD, Liccioli T, Schmid F, Kalatzis N, Sundstrom JF, Gardner JM, Jiranek V. 2014. Genome-wide identification of the Fermentome; genes required for successful and timely completion of wine-like fermentation by Saccharomyces cerevisiae. BMC genomics 15:552.

17. Cadière A, Aguera E, Caillé S, Ortiz-Julien A, Dequin S. 2012. Pilot-scale evaluation the enological traits of a novel, aromatic wine yeast strain obtained by adaptive evolution. Food microbiology 32:332–337.

18. Vila I. 1998. Les levures aromatiques en vinification: évaluation de ce caractère par l’analyse sensorielle et l’analyse chimique. Déterminisme biochimique des facteurs responsables. PhD Thesis, Montpellier 2.

19. Du Toit W, Marais J, Pretorius I, Du Toit M. 2006. Oxygen in must and wine: A review. South African Journal of Enology and Viticulture 27:76–94.

20. Roullier-Gall C, Boutegrabet L, Gougeon RD, Schmitt-Kopplin P. 2014. A grape and wine chemodiversity comparison of different appellations in Burgundy: Vintage vs terroir effects. Food chemistry 152:100–107.

21. Vilanova M, Rodríguez I, Canosa P, Otero I, Gamero E, Moreno D, Talaverano I, Valdés E. 2015. Variability in chemical composition of Vitis vinifera cv Mencía from different geographic areas and vintages in Ribeira Sacra (NW Spain). Food chemistry 169:187–196.

22. Ramos MC, Martínez de Toda F. 2019. Variability of Tempranillo grape composition in the Rioja DOCa (Spain) related to soil and climatic characteristics. Journal of the Science of Food and Agriculture 99:1153–1165.

23. Grainger C, Yeh A, Byer S, Hjelmeland A, Lima MM, Runnebaum RC. 2021. Vineyard site impact on the elemental composition of Pinot noir wines. Food Chemistry 334:127386.

24. Bokulich NA, Thorngate JH, Richardson PM, Mills DA. 2014. Microbial biogeography of wine grapes is conditioned by cultivar, vintage, and climate. Proceedings of the National Academy of Sciences 111:E139–E148.

25. Bokulich NA, Collins TS, Masarweh C, Allen G, Heymann H, Ebeler SE, Mills DA. 2016. Associations among wine grape microbiome, metabolome, and fermentation behavior suggest microbial contribution to regional wine characteristics. MBio 7.

26. David V, Terrat S, Herzine K, Claisse O, Rousseaux S, Tourdot-Maréchal R, Masneuf-Pomarede I, Ranjard L, Alexandre H. 2014. High-throughput sequencing of amplicons for monitoring yeast biodiversity in must and during alcoholic fermentation. Journal of industrial microbiology & biotechnology 41:811–821.

27. Pinto C, Pinho D, Cardoso R, Custódio V, Fernandes J, Sousa S, Pinheiro M, Egas C, Gomes AC. 2015. Wine fermentation microbiome: a landscape from different Portuguese wine appellations. Frontiers in microbiology 6:905.

28. Wang C, García-Fernández D, Mas A, Esteve-Zarzoso B. 2015. Fungal diversity in grape must and wine fermentation assessed by massive sequencing, quantitative PCR and DGGE. Frontiers in microbiology 6:1156.

29. Garofalo C, Russo P, Beneduce L, Massa S, Spano G, Capozzi V. 2016. Non-Saccharomyces biodiversity in wine and the ‘microbial terroir’: A survey on Nero di Troia wine from the Apulian region, Italy. Annals of microbiology 66:143–150.

30. Mezzasalma V, Sandionigi A, Bruni I, Bruno A, Lovicu G, Casiraghi M, Labra M. 2017. Grape microbiome as a reliable and persistent signature of field origin and environmental conditions in Cannonau wine production. PLoS One 12:e0184615.

31. Mezzasalma V, Sandionigi A, Guzzetti L, Galimberti A, Grando MS, Tardaguila J, Labra M. 2018. Geographical and cultivar features differentiate grape microbiota in Northern Italy and Spain vineyards. Frontiers in microbiology 9:946.

32. Singh P, Santoni S, This P, Péros J-P. 2018. Genotype-environment interaction shapes the microbial assemblage in grapevine’s phyllosphere and carposphere: an NGS approach. Microorganisms 6:96.

33. 33. Liu D, Chen Q, Zhang P, Chen D, Howell KS. 2020. The Fungal Microbiome Is an Important Component of Vineyard Ecosystems and Correlates with Regional Distinctiveness of Wine. mSphere 5.

34. Jolly N, Augustyn O, Pretorius I. 2003. The occurrence of non-Saccharomyces cerevisiae yeast species over three vintages in four vineyards and grape musts from four production regions of the Western Cape, South Africa. South African Journal of Enology and Viticulture 24:35–42.

35. Ghosh S, Bagheri B, Morgan HH, Divol B, Setati ME. 2015. Assessment of wine microbial diversity using ARISA and cultivation-based methods. Annals of microbiology 65:1833–1840.

36. Wang C, Esteve-Zarzoso B, Cocolin L, Mas A, Rantsiou K. 2015. Viable and culturable populations of Saccharomyces cerevisiae, Hanseniaspora uvarum and Starmerella bacillaris (synonym Candida zemplinina) during Barbera must fermentation. Food Research International 78:195–200.

37. Bagheri B, Bauer F, Setati M. 2015. The diversity and dynamics of indigenous yeast communities in grape must from vineyards employing different agronomic practices and their influence on wine fermentation. South African Journal of Enology and Viticulture 36:243–251.

38. Bagheri B, Bauer FF, Cardinali G, Setati ME. 2020. ecological interactions are a primary driver of population dynamics in wine yeast microbiota during fermentation. Scientific reports 10:1–12.

39. Brou P, Taillandier P, Beaufort S, Brandam C. 2018. Mixed culture fermentation using Torulaspora delbrueckii and Saccharomyces cerevisiae with direct and indirect contact: impact of anaerobic growth factors. European Food Research and Technology 244:1699–1710.

40. Curiel JA, Morales P, Gonzalez R, Tronchoni J. 2017. Different non-Saccharomyces yeast species stimulate nutrient consumption in S. cerevisiae mixed cultures. Frontiers in Microbiology 8:2121.

41. Alonso-del-Real J, Pérez-Torrado R, Querol A, Barrio E. 2019. Dominance of wine Saccharomyces cerevisiae strains over S. kudriavzevii in industrial fermentation competitions is related to an acceleration of nutrient uptake and utilization. Environmental microbiology 21:1627–1644.

42. Tronchoni J, Curiel JA, Morales P, Torres-Pérez R, Gonzalez R. 2017. Early transcriptional response to biotic stress in mixed starter fermentations involving Saccharomyces cerevisiae and Torulaspora delbrueckii. International journal of food microbiology 241:60–68.

43. Shekhawat K, Patterton H, Bauer FF, Setati ME. 2019. RNA-seq based transcriptional analysis of Saccharomyces cerevisiae and Lachancea thermotolerans in mixed-culture fermentations under anaerobic conditions. BMC genomics 20:145.

44. Conacher CG, Rossouw D, Bauer F. 2019. Peer pressure: evolutionary responses to biotic pressures in wine yeasts. FEMS Yeast Research 19:foz072.

45. Bordet F, Joran A, Klein G, Roullier-Gall C, Alexandre H. 2020. Yeast–Yeast Interactions: Mechanisms, Methodologies and Impact on Composition. Microorganisms 8:600.

46. Egli C, Edinger W, Mitrakul C, Henick-Kling T. 1998. Dynamics of indigenous and inoculated yeast populations and their effect on the sensory character of Riesling and Chardonnay wines. Journal of Applied Microbiology 85:779–789.

47. Bartowsky EJ. 2009. Bacterial spoilage of wine and approaches to minimize it. Letters in Applied Microbiology 48:149–156.

48. Cantu A, Lafontaine S, Frias I, Sokolowsky M, Yeh A, Lestringant P, Hjelmeland A, Byer S, Heymann H, Runnebaum RC. 2021. Investigating the impact of regionality on the sensorial and chemical aging characteristics of Pinot noir grown throughout the US West coast. Food Chemistry 337:127720.

49. Bisson LF. 2019. Gene Expression in Yeasts During Wine Fermentation, p. 165–209. In Yeasts in the Production of Wine. Springer.

50. Rodicio R, Heinisch JJ. 2017. Carbohydrate Metabolism in Wine Yeasts, p. 189–213. In Biology of Microorganisms on Grapes, in Must and in Wine. Springer International Publishing.

51. Ozcan S, Johnston M. 1996. Two different repressors collaborate to restrict expression of the yeast glucose transporter genes HXT2 and HXT4 to low levels of glucose. Molecular and Cellular Biology 16:5536–5545.

52. Ogawa N, DeRisi J, Brown PO. 2000. New Components of a System for Phosphate Accumulation and Polyphosphate Metabolism inSaccharomyces cerevisiaeRevealed by Genomic Expression Analysis. Molecular Biology of the Cell 11:4309–4321.

53. Ljungdahl PO, Daignan-Fornier B. 2012. Regulation of amino acid, nucleotide, and phosphate metabolism in Saccharomyces cerevisiae. Genetics 190:885–929.

54. Barbosa C, Mendes-Faia A, Lage P, Mira NP, Mendes-Ferreira A. 2015. Genomic expression program of Saccharomyces cerevisiae along a mixed-culture wine fermentation with Hanseniaspora guilliermondii. Microbial cell factories 14:124.

55. Kosel J, Čadež N, Schuller D, Carreto L, Franco-Duarte R, Raspor P. 2017. The influence of Dekkera bruxellensis on the transcriptome of Saccharomyces cerevisiae and on the aromatic profile of synthetic wine must. FEMS Yeast Research 17:fox018.

56. Reiter T, Montpetit R, Byer S, Frias I, Leon E, Viano R, Mcloughlin M, Halligan T, Hernandez D, Figueroa-Balderas R, Cantu D, Steenwerth K, Runnebaum R, Montpetit B. 2021. Transcriptomics provides a genetic signature of vineyard site with insight into vintage-independent regional wine characteristics. bioRxiv 2021.01.07.425830.

57. Devatine A, Chiciuc I, Mietton-Peuchot M. 2011. The protective role of dissolved carbon dioxide against wine oxidation: a simple and rational approach. OENO One 45:189–197.

58. Moenne MI, Saa P, Laurie VF, Pérez-Correa JR, Agosin E. 2014. Oxygen incorporation and dissolution during industrial-scale red wine fermentations. Food and bioprocess technology 7:2627–2636.

59. Abramova N, Sertil O, Mehta S, Lowry CV. 2001. Reciprocal regulation of anaerobic and aerobic cell wall mannoprotein gene expression in Saccharomyces cerevisiae. Journal of bacteriology 183:2881–2887.

60. Sertil O, Kapoor R, Cohen BD, Abramova N, Lowry CV. 2003. Synergistic repression of anaerobic genes by Mot3 and Rox1 in Saccharomyces cerevisiae. Nucleic acids research 31:5831–5837.

61. Kwast KE, Burke PV, Poyton RO. 1998. Oxygen sensing and the transcriptional regulation of oxygen-responsive genes in yeast. Journal of Experimental Biology 201:1177–1195.

62. Wöhl T, Klier H, Ammer H, Lottspeich F, Magdolen V. 1993. The HYP2 gene of Saccharomyces cerevisiae is essential for aerobic growth: characterization of different isoforms of the hypusine-containing protein Hyp2p and analysis of gene disruption mutants. Molecular and General Genetics MGG 241–241:305–311.

63. Burke PV, Poyton RO. 1998. Structure/function of oxygen-regulated isoforms in cytochrome c oxidase. Journal of Experimental Biology 201:1163–1175.

64. Remize F, Cambon B, Barnavon L, Dequin S. 2003. Glycerol formation during wine fermentation is mainly linked to Gpd1p and is only partially controlled by the HOG pathway. Yeast 20:1243–1253.

65. Jung J-Y, Kim T-Y, Ng C-Y, Oh M-K. 2012. Characterization of GCY1inSaccharomyces cerevisiaeby metabolic profiling. Journal of Applied Microbiology 113:1468–1478.

66. Kliewer WM. 1970. Free Amino Acids and Other Nitrogenous Fractions in Wine Grapes. Journal of Food Science 35:17–21.

67. Schure EG ter, Riel NAW van, Verrips CT. 2000. The role of ammonia metabolism in nitrogen catabolite repression inSaccharomyces cerevisiae. FEMS Microbiology Reviews 24:67–83.

68. Huang HL, Brandriss MC. 2000. The Regulator of the Yeast Proline Utilization Pathway Is Differentially Phosphorylated in Response to the Quality of the Nitrogen Source. Molecular and Cellular Biology 20:892–899.

69. Takagi H, Taguchi J, Kaino T. 2016. Proline accumulation protectsSaccharomyces cerevisiaecells in stationary phase from ethanol stress by reducing reactive oxygen species levels. Yeast 33:355–363.

70. Rosenfeld E, Beauvoit B, Blondin B, Salmon J-M. 2003. Oxygen consumption by anaerobic Saccharomyces cerevisiae under enological conditions: effect on fermentation kinetics. Applied and environmental microbiology 69:113–121.

71. Tarko T, Duda-Chodak A, Sroka P, Siuta M. 2020. The Impact of Oxygen at Various Stages of Vinification on the Chemical Composition and the Antioxidant and Sensory Properties of White and Red Wines. International Journal of Food Science 2020.

72. O’Connor-Cox E, Lodolo E, Axcell B. 1996. Mitochondrial relevance to yeast fermentative performance: a review. Journal of the Institute of Brewing 102:19–25.

73. Kitagaki H, Takagi H. 2014. Mitochondrial metabolism and stress response of yeast: applications in fermentation technologies. Journal of bioscience and bioengineering 117:383–393.

74. Zara G, van Vuuren HJ, Mannazzu I, Zara S, Budroni M. 2019. Transcriptomic response of Saccharomyces cerevisiae during fermentation under oleic acid and ergosterol depletion. Fermentation 5:57.

75. Zeng J, Smith KE, Chong P. 1993. Effects of alcohol-induced lipid interdigitation on proton permeability in L-alpha-dipalmitoylphosphatidylcholine vesicles. Biophysical journal 65:1404–1414.

76. Landolfo S, Politi H, Angelozzi D, Mannazzu I. 2008. ROS accumulation and oxidative damage to cell structures in Saccharomyces cerevisiae wine strains during fermentation of high-sugar-containing medium. Biochimica et Biophysica Acta (BBA)-General Subjects 1780:892–898.

77. Molenaar D, Van Berlo R, De Ridder D, Teusink B. 2009. Shifts in growth strategies reflect tradeoffs in cellular economics. Molecular systems biology 5:323.

78. Wimpenny J. 1969. The effect of Eh on regulatory processes in facultative anaerobes. Biotechnology and bioengineering 11:623–629.

79. Somlo M, Fukuhara H. 1965. On the necessity of molecular oxygen for the synthesis of respiratory enzymes in yeast. Biochemical and Biophysical Research Communications 19:587–591.

80. Pammer M, Briza P, Ellinger A, Schuster T, Stucka R, Feldmann H, Breitenbach M. 1992. DIT101 (CSD2, CAL1), a cell cycle-regulated yeast gene required for synthesis of chitin in cell walls and chitosan in spore walls. Yeast 8:1089–1099.

81. Argüello-Miranda O, Liu Y, Wood NE, Kositangool P, Doncic A. 2018. Integration of multiple metabolic signals determines cell fate prior to commitment. Molecular cell 71:733–744.

82. Zhao H, Wang Q, Liu C, Shang Y, Wen F, Wang F, Liu W, Xiao W, Li W. 2018. A role for the respiratory chain in regulating meiosis initiation in Saccharomyces cerevisiae. Genetics 208:1181–1194.

83. Sipiczki M. 2011. Diversity, variability and fast adaptive evolution of the wine yeast (Saccharomyces cerevisiae) genome—a review. Annals of microbiology 61:85–93.

84. Reiner S, Micolod D, Zellnig G, Schneiter R. 2006. A genomewide screen reveals a role of mitochondria in anaerobic uptake of sterols in yeast. Molecular Biology of the Cell 17:90–103.

85. Perez-Gallardo RV, Briones LS, Díaz-Pérez AL, Gutiérrez S, Rodríguez-Zavala JS, Campos-García J. 2013. Reactive oxygen species production induced by ethanol in Saccharomyces cerevisiae increases because of a dysfunctional mitochondrial iron– sulfur cluster assembly system. FEMS yeast research 13:804–819.

86. Carmel-Harel O, Storz G. 2000. Roles of the glutathione-and thioredoxin-dependent reduction systems in the Escherichia coli and Saccharomyces cerevisiae responses to oxidative stress. Annual Reviews in Microbiology 54:439–461.

87. Toledano MB, Delaunay-Moisan A, Outten CE, Igbaria A. 2013. Functions and cellular compartmentation of the thioredoxin and glutathione pathways in yeast. Antioxidants & redox signaling 18:1699–1711.

88. Bridi R, González A, Bordeu E, López-Alarcón C, Aspée A, Diethelm B, Lissi E, Parpinello GP, Versari A. 2015. Monitoring peroxides generation during model wine fermentation by FOX-1 assay. Food chemistry 175:25–28.

89. Maslanka R, Zadrag-Tecza R, Kwolek-Mirek M. 2020. Linkage between Carbon Metabolism, Redox Status and Cellular Physiology in the Yeast Saccharomyces cerevisiae Devoid of SOD1 or SOD2 Gene. Genes 11:780.

90. Matsufuji Y, Yamamoto K, Yamauchi K, Mitsunaga T, Hayakawa T, Nakagawa T. 2013. Novel physiological roles for glutathione in sequestering acetaldehyde to confer acetaldehyde tolerance in Saccharomyces cerevisiae. Applied microbiology and biotechnology 97:297–303.

91. Koc A, Mathews CK, Wheeler LJ, Gross MK, Merrill GF. 2006. Thioredoxin is required for deoxyribonucleotide pool maintenance during S phase. Journal of Biological Chemistry 281:15058–15063.

92. Muller E. 1991. Thioredoxin deficiency in yeast prolongs S phase and shortens the G1 interval of the cell cycle. Journal of Biological Chemistry 266:9194–9202.

93. Shenton D, Grant CM. 2003. Protein S-thiolation targets glycolysis and protein synthesis in response to oxidative stress in the yeast Saccharomyces cerevisiae. Biochemical Journal 374:513–519.

94. Ralser M, Wamelink MM, Kowald A, Gerisch B, Heeren G, Struys EA, Klipp E, Jakobs C, Breitenbach M, Lehrach H, others. 2007. Dynamic rerouting of the carbohydrate flux is key to counteracting oxidative stress. Journal of biology 6:10.

95. Deponte M. 2017. The incomplete glutathione puzzle: just guessing at numbers and figures? Antioxidants & redox signaling 27:1130–1161.

96. Saint-Prix F, Bönquist L, Dequin S. 2004. Functional analysis of the ALD gene family of Saccharomyces cerevisiae during anaerobic growth on glucose: the NADP+-dependent Ald6p and Ald5p isoforms play a major role in acetate formation. Microbiology 150:2209–2220.

97. Greetham D, Vickerstaff J, Shenton D, Perrone GG, Dawes IW, Grant CM. 2010. Thioredoxins function as deglutathionylase enzymes in the yeast Saccharomyces cerevisiae. BMC biochemistry 11:3.

98. Collinson EJ, Wheeler GL, Garrido EO, Avery AM, Avery SV, Grant CM. 2002. The yeast glutaredoxins are active as glutathione peroxidases. Journal of Biological Chemistry 277:16712–16717.

99. Ohdate T, Kita K, Inoue Y. 2010. Kinetics and redox regulation of Gpx1, an atypical 2-Cys peroxiredoxin, in Saccharomyces cerevisiae. FEMS yeast research 10:787–790.

100. Ohdate T, Izawa S, Kita K, Inoue Y. 2010. Regulatory mechanism for expression of GPX1 in response to glucose starvation and Ca2+ in Saccharomyces cerevisiae: involvement of Snf1 and Ras/cAMP pathway in Ca2+ signaling. Genes to Cells 15:59–75.

101. Greetham D, Grant CM. 2009. Antioxidant activity of the yeast mitochondrial one-Cys peroxiredoxin is dependent on thioredoxin reductase and glutathione in vivo. Molecular and cellular biology 29:3229–3240.

102. Martins AMT, Cordeiro CAA, Freire AMJP. 2001. In situ analysis of methylglyoxal metabolism in Saccharomyces cerevisiae. FEBS letters 499:41–44.

103. Dhaoui M, Auchère F, Blaiseau P-L, Lesuisse E, Landoulsi A, Camadro J-M, Haguenauer-Tsapis R, Belgareh-Touzé N. 2011. Gex1 is a yeast glutathione exchanger that interferes with pH and redox homeostasis. Molecular biology of the cell 22:2054– 2067.

104. Cordente AG, Capone DL, Curtin CD. 2015. Unravelling glutathione conjugate catabolism in Saccharomyces cerevisiae: the role of glutathione/dipeptide transporters and vacuolar function in the release of volatile sulfur compounds 3-mercaptohexan-1-ol and 4-mercapto-4-methylpentan-2-one. Applied microbiology and biotechnology 99:9709–9722.

105. Kumar A, Tikoo S, Maity S, Sengupta S, Sengupta S, Kaur A, Kumar Bachhawat A. 2012. Mammalian proapoptotic factor ChaC1 and its homologues function as γ-glutamyl cyclotransferases acting specifically on glutathione. EMBO reports 13:1095–1101.

106. Chen X, Li S, Liu L. 2014. Engineering redox balance through cofactor systems. Trends in biotechnology 32:337–343.

107. Fariña L, Medina K, Urruty M, Boido E, Dellacassa E, Carrau F. 2012. Redox effect on volatile compound formation in wine during fermentation by Saccharomyces cerevisiae. Food chemistry 134:933–939.

108. Bloem A, Sanchez I, Dequin S, Camarasa C. 2016. Metabolic impact of redox cofactor perturbations on the formation of aroma compounds in Saccharomyces cerevisiae. Applied and environmental microbiology 82:174–183.

109. Xu X, Bao M, Niu C, Wang J, Liu C, Zheng F, Li Y, Li Q. 2019. Engineering the cytosolic NADH availability in lager yeast to improve the aroma profile of beer. Biotechnology letters 41:363–369.

110. Xu X, Song Y, Guo L, Cheng W, Niu C, Wang J, Liu C, Zheng F, Zhou Y, Li X, others. 2019. Higher NADH Availability of Lager Yeast Increases the Flavor Stability of Beer. Journal of Agricultural and Food Chemistry 68:584–590.

111. Causton HC, Ren B, Koh SS, Harbison CT, Kanin E, Jennings EG, Lee TI, True HL, Lander ES, Young RA. 2001. Remodeling of yeast genome expression in response to environmental changes. Molecular biology of the cell 12:323–337.

112. Mbuyane LL, de Kock M, Bauer FF, Divol B. 2018. Torulaspora delbrueckii produces high levels of C5 and C6 polyols during wine fermentations. FEMS yeast research 18:foy084.

113. Endo A, Futagawa-Endo Y, Dicks LM. 2009. Isolation and characterization of fructophilic lactic acid bacteria from fructose-rich niches. Systematic and Applied Microbiology 32:593–600.

114. Du Toit M, Pretorius IS. 2000. Microbial spoilage and preservation of wine: using weapons for nature’s own arsenal.

115. Quain DE, Boulton CA. 1987. Growth and Metabolism of Mannitol by Strains of Saccharomyces cerevisiae. Microbiology 133:1675–1684.

116. Jordan P, Choe J-Y, Boles E, Oreb M. 2016. Hxt13, Hxt15, Hxt16 and Hxt17 from Saccharomyces cerevisiae represent a novel type of polyol transporters. Scientific Reports 6.

117. Ramakrishnan V, Walker GA, Fan Q, Ogawa M, Luo Y, Luong P, Joseph C, Bisson LF. 2016. Inter-kingdom modification of metabolic behavior:[GAR+] prion induction in Saccharomyces cerevisiae mediated by wine ecosystem bacteria. Frontiers in Ecology and Evolution 4:137.

118. Brown JC, Lindquist S. 2009. A heritable switch in carbon source utilization driven by an unusual yeast prion. Genes & development 23:2320–2332.

119. Jarosz DF, Brown JC, Walker GA, Datta MS, Ung WL, Lancaster AK, Rotem A, Chang A, Newby GA, Weitz DA, others. 2014. Cross-kingdom chemical communication drives a heritable, mutually beneficial prion-based transformation of metabolism. Cell 158:1083–1093.

120. Nurgel C, Pickering G. 2005. Contribution of glycerol, ethanol and sugar to the perception of viscosity and density elicited by model white wines. Journal of texture studies 36:303–323.

121. Ansell R, Granath K, Hohmann S, Thevelein JM, Adler L. 1997. The two isoenzymes for yeast NAD+-dependent glycerol 3-phosphate dehydrogenase encoded by GPD1 and GPD2 have distinct roles in osmoadaptation and redox regulation. The EMBO journal 16:2179–2187.

122. Remize F, Barnavon L, Dequin S. 2001. Glycerol export and glycerol-3-phosphate dehydrogenase, but not glycerol phosphatase, are rate limiting for glycerol production in Saccharomyces cerevisiae. Metabolic engineering 3:301–312.

123. Smith KD, Gordon PB, Rivetta A, Allen KE, Berbasova T, Slayman C, Strobel SA. 2015. Yeast Fex1p Is a Constitutively Expressed Fluoride Channel with Functional Asymmetry of Its Two Homologous Domains. Journal of Biological Chemistry 290:19874–19887.

124. Clayton MG. 1997. Fluoride inhibition of wine yeasts: a thesis presented in partial fulfilment of the requirements for the degress of Master of Science in Microbiology at Massey University. PhD Thesis, Massey University.

125. Dobin A, Davis CA, Schlesinger F, Drenkow J, Zaleski C, Jha S, Batut P, Chaisson M, Gingeras TR. 2013. STAR: ultrafast universal RNA-seq aligner. Bioinformatics 29:15–21.

126. Smith T, Heger A, Sudbery I. 2017. UMI-tools: modeling sequencing errors in Unique Molecular Identifiers to improve quantification accuracy. Genome research 27:491–499.

127. Anders S, Pyl PT, Huber W. 2015. HTSeq—a Python framework to work with high-throughput sequencing data. Bioinformatics 31:166–169.

128. Robinson MD, McCarthy DJ, Smyth GK. 2010. edgeR: a Bioconductor package for differential expression analysis of digital gene expression data. Bioinformatics 26:139– 140.

129. Ritchie ME, Phipson B, Wu D, Hu Y, Law CW, Shi W, Smyth GK. 2015. limma powers differential expression analyses for RNA-sequencing and microarray studies. Nucleic acids research 43:e47–e47.

130. Liebermeister W, Noor E, Flamholz A, Davidi D, Bernhardt J, Milo R. 2014. Visual account of protein investment in cellular functions. Proceedings of the National Academy of Sciences 111:8488–8493.

131. Yu G, Wang L-G, Han Y, He Q-Y. 2012. clusterProfiler: an R package for comparing biological themes among gene clusters. Omics: a journal of integrative biology 16:284– 287.

132. Rorbach J, Bobrowicz A, Pearce S, Minczuk M. 2014. Polyadenylation in bacteria and organelles, p. 211–227. In Polyadenylation. Springer.

133. Brown CT, Irber L. 2016. sourmash: a library for MinHash sketching of DNA. Journal of Open Source Software 1:27.

134. Pierce NT, Irber L, Reiter T, Brooks P, Brown CT. 2019. Large-scale sequence comparisons with sourmash. F1000Research 8.

135. Li H. 2013. Aligning sequence reads, clone sequences and assembly contigs with BWA-MEM. arXiv preprint arXiv:13033997.

136. Ciriacy M. 1975. Genetics of alcohol dehydrogenase inSaccharomyces cerevisiac. Molecular and General Genetics MGG 138:157–164.

137. Zuzuarregui A, Monteoliva L, Gil C, others. 2006. Transcriptomic and proteomic approach for understanding the molecular basis of adaptation of Saccharomyces cerevisiae to wine fermentation. Applied and environmental microbiology 72:836–847.

138. Curiel JA, Salvadó Z, Tronchoni J, Morales P, Rodrigues AJ, Quirós M, Gonzalez R. 2016. Identification of target genes to control acetate yield during aerobic fermentation with Saccharomyces cerevisiae. Microbial cell factories 15:156.

139. Piper P, Mahé Y, Thompson S, Pandjaitan R, Holyoak C, Egner R, Mühlbauer M, Coote P, Kuchler K. 1998. The Pdr12 ABC transporter is required for the development of weak organic acid resistance in yeast. The EMBO journal 17:4257–4265.

140. Caro LHP, Smits GJ, van Egmond P, Chapman JW, Klis FM. 1998. Transcription of multiple cell wall protein-encoding genes in Saccharomyces cerevisiae is differentially regulated during the cell cycle. FEMS microbiology letters 161:345–349.

141. Crépin L, Truong NM, Bloem A, Sanchez I, Dequin S, Camarasa C. 2017. Management of multiple nitrogen sources during wine fermentation by Saccharomyces cerevisiae. Applied and environmental microbiology 83.

142. Hazelwood LA, Daran J-M, Van Maris AJ, Pronk JT, Dickinson JR. 2008. The Ehrlich pathway for fusel alcohol production: a century of research on Saccharomyces cerevisiae metabolism. Applied and environmental microbiology 74:2259–2266.

143. Cebollero E, Reggiori F. 2009. Regulation of autophagy in yeast Saccharomyces cerevisiae. Biochimica et Biophysica Acta (BBA) - Molecular Cell Research 1793:1413– 1421.

144. Alexandre H, Ansanay-Galeote V, Dequin S, Blondin B. 2001. Global gene expression during short-term ethanol stress inSaccharomyces cerevisiae. FEBS Letters 498:98–103.

145. Lillie SH, Pringle JR. 1980. Reserve carbohydrate metabolism in Saccharomyces cerevisiae: responses to nutrient limitation. Journal of Bacteriology 143:1384–1394.

146. François J, Parrou JL. 2001. Reserve carbohydrates metabolism in the yeastSaccharomyces cerevisiae. FEMS Microbiology Reviews 25:125–145.

147. Singer MA, Lindquist S. 1998. Multiple Effects of Trehalose on Protein Folding In Vitro and In Vivo. Molecular Cell 1:639–648.

148. Parrou JL, Teste M-A, Francois J. 1997. Effects of various types of stress on the metabolism of reserve carbohydrates in Saccharomyces cerevisiae: genetic evidence for a stress-induced recycling of glycogen and trehalose. Microbiology 143:1891–1900.

149. Udom N, Chansongkrow P, Charoensawan V, Auesukaree C. 2019. Coordination of the Cell Wall Integrity and High-Osmolarity Glycerol Pathways in Response to Ethanol Stress in Saccharomyces cerevisiae. Applied and Environmental Microbiology 85.

